# Effects of climate warming on the pine processionary moth at the southern edge of its range: a retrospective analysis on egg survival in Tunisia

**DOI:** 10.1101/2021.08.17.456665

**Authors:** Asma Bourougaaoui, Christelle Robinet, Mohamed L. Ben Jamaa, Mathieu Laparie

## Abstract

In recent years, ectotherm species have largely been impacted by extreme climate events, essentially heatwaves. In Tunisia, the pine processionary moth (PPM), *Thaumetopoea pityocampa*, is a highly damaging pine defoliator, which typically lays eggs in summer. Its geographical range is expanding northwards in Europe while retracting from South Tunisia where summer temperatures can reach extremely high values. In this study, we aimed at exploring the effects of climate change on this species at its southern range edge. We investigated variations of fecundity and causes of egg mortality over time using historical and contemporary collections of egg masses from different Tunisian sites to seek relationships with regional climate change over three decades (1990-2019). Our results suggest negative effects of summer heat on egg survival, reflected in a decrease of hatching rate down to 0% in one site during a heatwave. Such a high hatching failure was found to result from both high egg sterility (our results did not allow distinguishing impeded mating success from failed egg maturation or early death of the embryo) and increased abortion of more developed embryos, but little effects of parasitism rate, thereby suggesting vulnerability to heat during embryonic development. We also observed decreasing female fecundity (*i.e*., number of eggs laid per female) in regions where data were available both in the 1990s and the 2010s, which was associated with a decrease in parasitism rate, while the climatic variability increased. This study investigated direct hatching failure in nature that may be related to the magnitude of warming in summer. Previous studies have confirmed the thermal sensitivity of early instars of the PPM to temperatures observed in the present work, including one population from South Tunisia. However, further work is required to evaluate the relative importance of warming summers among populations because the risk of heat stress depends on the phenology of sensitive instars, and populations from the warmest areas may not necessarily be the most vulnerable to climate change if they already evolved phenological heat avoidance. In addition to heat-induced mortality, the ultimate fitness of individuals that survive challenging heat stresses during early developmental stages should also be explored to determine potential carry-over effects on subsequent life stages.

## Introduction

During the period 1901-2010, land temperature has risen by 1.12 and 0.84°C in the Northern and Southern hemispheres, respectively (Jones et al., 2012). Climate change is predicted to increase not only mean temperatures but also temperature variability and, in turn, the magnitude and frequency of stochastic extreme thermal events (Allen et al., 2012). This is already being increasingly observed over most parts of the world (Allen et al., 2012; Coumou & Rahmstorf, 2012; Fischer & Schär, 2010), particularly northern Africa (Fontaine et al., 2013; Nangombe et al., 2019; Zittis et al., 2021). Mean temperature has risen by about 1.4°C since 1901 in Tunisia, with a remarkable average increase of +0.4°C per decade in the last 30 years, primarily observed during summer in southern regions where temperatures can exceed 40°C (Verner et al., 2013). Together with average warming, increasing thermal fluctuations and extreme events may impact all fitness components (*e.g*., phenology, morphology, behaviour, locomotor activity, and physiology) of organisms (Charmantier & Gienapp, 2014; Chuine, 2010; Chuine et al., 2013; Gardner et al., 2011; Ghosh et al., 2013; Kingsolver et al., 2013; Liu et al., 1995; Pigliucci, 2001, 2005; Pincebourde et al., 2021; Pincebourde & Woods, 2020; Sheridan & Bickford, 2011; Thompson et al., 2013; Woods et al., 2015; Wu et al., 2019). Moreover, the concomitant and mutually interacting facets of climate change may ultimately translate into survival and in turn alter genetic frequencies, population density in given habitats, as well as persistence and distribution of many organisms (Root et al., 2003; Vasseur et al., 2014). In the twentieth century, a wide range of taxa ranging from invertebrates to mammals and from grasses to trees have shifted their ranges poleward, upslope or both (Crozier, 2004; Hickling et al., 2005; Karban & Strauss, 2004; Parmesan et al., 1999; Parmesan & Yohe, 2003; Root et al., 2003; Walther et al., 2002).

The pine processionary moth (hereafter referred to as PPM), *Thaumetopoea pityocampa* (Denis & Schiffermüller, 1776) (Lepidoptera, Notodontidae), is a highly damaging pest of pine forests across the circum-Mediterranean region (Carus, 2009; Démolin, 1969; Jacquet et al., 2013; Sbay & Zas, 2018). The geographic range of the PPM extends from northern Africa to southern Europe, from the Atlantic coast to the western part of Turkey (EPPO, 2004; Roques, 2015). The PPM is a well-documented insect that has been acknowledged by the Intergovernmental Panel on Climate Change (IPCC) as one of the few species for which the causal relationship between climate warming and range expansion has been thoroughly proven (Battisti et al., 2005; Rosenzweig et al., 2007). The distribution range remained relatively steady until the late 1990s but then expanded towards higher latitudes and elevations in southern Europe. Indeed, warming winter temperatures have facilitated feeding in this winter-developing species and thus indirectly contributed to improving survival rate and growth rate in newly colonized areas (Battisti et al., 2005, Robinet et al., 2007).

Contrary to the beneficial effects of climate change demonstrated near the northern distribution edge of the PPM, adverse effects of climate change have been observed on the southern range edge (North Africa). Range retraction has been described in southern Tunisia and was found to result from increasing mortality rates of early life stages in a translocation experiment along a natural thermal gradient, which could be ascribed to local effects of climate warming (Bourougaaoui et al., 2021). Fecundity, hatching rate and predation at the egg stage (mostly from parasitoids) presumably play an important role in the PPM because this species is gregarious. Several studies have emphasized how larval performance depends on the realised group size, i.e. the number of neonates, and ultimately the survival of the whole colony until the end of larval growth (Clark & Faeth, 1997; Denno & Benrey, 1997; Ronnås et al., 2010). Colony density has been suggested to influence feeding activity and feeding efficiency of individual larvae, which is particularly critical in early stages when individuals have little desiccation and starvation resistances. The number of larvae was also shown to impact silk weaving activity to build and maintain the nests that shelter larvae during the day until their pupation in spring (Démolin, 1965; Martin, 2005). As a result, the number of surviving tents and the average proportion of living larvae per tent were positively correlated to colony size (Pérez-Contreras et al., 2003; Roques et al., 2015). Focusing on the main drivers of colony size in early development is therefore of key importance to understand distribution changes and responses to climate change.

In Tunisia, the life cycle of PPM is generally univoltine, however it can extend over two years at high altitudes due to prolonged diapause in a fraction of the pupae (Roques, 2015). Flight periods are poorly documented, nonetheless a study conducted by Démolin and Rive in 1968 in high and medium latitudes, revealed that most individuals fly in the second half of July at high elevations and August to September at mid elevations (Ben Jamâa & Jerraya, 1999; Démolin & Rive, 1968). Due to the short lifespan of adults, egg laying occurs immediately after adult flights, and eggs and neonate larvae are presumably the instars that are most likely exposed to acute heat during the whole life cycle. Understanding the effects of warming on female fecundity, egg survival and egg parasitoids is crucial to explore the overall effects of climate change of this species at its southern range edge where warming is known to be of great magnitude.

In this study, we explored how climate warming over the last three decades may have impacted egg survival and hatching rate in Tunisia. To address this question, we combined historical and contemporary collections of egg masses originating from different Tunisian localities in the 1990s (1992, 1993, and 1995) and in the 2010s (2010, 2014, 2017, 2018, and 2019). Egg phenotypes and survival rate were investigated with regard to regional climatic features and contrasts analyzed from 30-year climatic data series across Tunisia. A cornerstone of this study is the identification of climate regions computed from multiple meteorological series, which allows comparing eggs from multiple sites within statistically consistent climates, instead of using arbitrary groups such as administrative regions. A grouping method was mandatory to analyze the long term data available on PPM eggs because exact sampling sites have changed over the years.

## Materials and Methods

### 1 Historical data (1992-2014) and egg sampling done for this study (2017-2019)

A total of 755 egg masses from historical datasets and recent collections were analyzed in this study. Egg masses originated from 22 sites distributed across the PPM distribution in Tunisia (Table 1; Figure 1; Table SM1). Historical datasets on egg masses collected in 1992, 1993, 1995, 2010 and 2014 on Aleppo pine stands, *Pinus halepensis* Miller, were retrieved from institutional reports (unpublished data, INRGREF). These datasets report the length of egg masses, the number of eggs per egg mass, and the phenotype of individual eggs (parasitized, aborted, sterile, hatched). In addition, we collected egg masses in various locations in 2017, 2018, and 2019. All these egg masses were also collected on Aleppo pine stands, before hatching but as late as possible in each region to ensure eggs were exposed to natural conditions, and then and kept at ambient temperature (25 ± 2°C) at the INRGREF laboratory near Tunis where we followed a protocol similar to that used for historical collections of egg masses. Egg masses were kept individually in test tubes capped with cotton to allow ventilation. Egg hatching was checked daily. After a period of at least 40 days with no additional hatching, the protective scales that cover PPM egg masses were removed to observe individual eggs under a binocular magnifier and collect data similar to that available in historical datasets: length of egg masses, number of eggs per egg mass (fecundity), and egg phenotype. First, hatched eggs were distinguished from unhatched eggs based on the presence of the characteristic large jagged exit hole from which the neonate left the egg, and an empty transparent shell. Then, unhatched eggs were dissected to assess the cause of mortality (parasitized, *i.e*., eggs with a small parasitoid exit hole and/or containing a dead parasitoid and/or containing parasitoid meconium; aborted, *i.e*., dead embryo or dead PPM larva; and sterile, *i.e*., undeveloped egg with dried-up yolk) (Imbert, 2012). Parasitism rate was calculated taking into account both emerged parasitoids found in the test tubes and dead ones found inside unhatched eggs.

**Figure 1.**
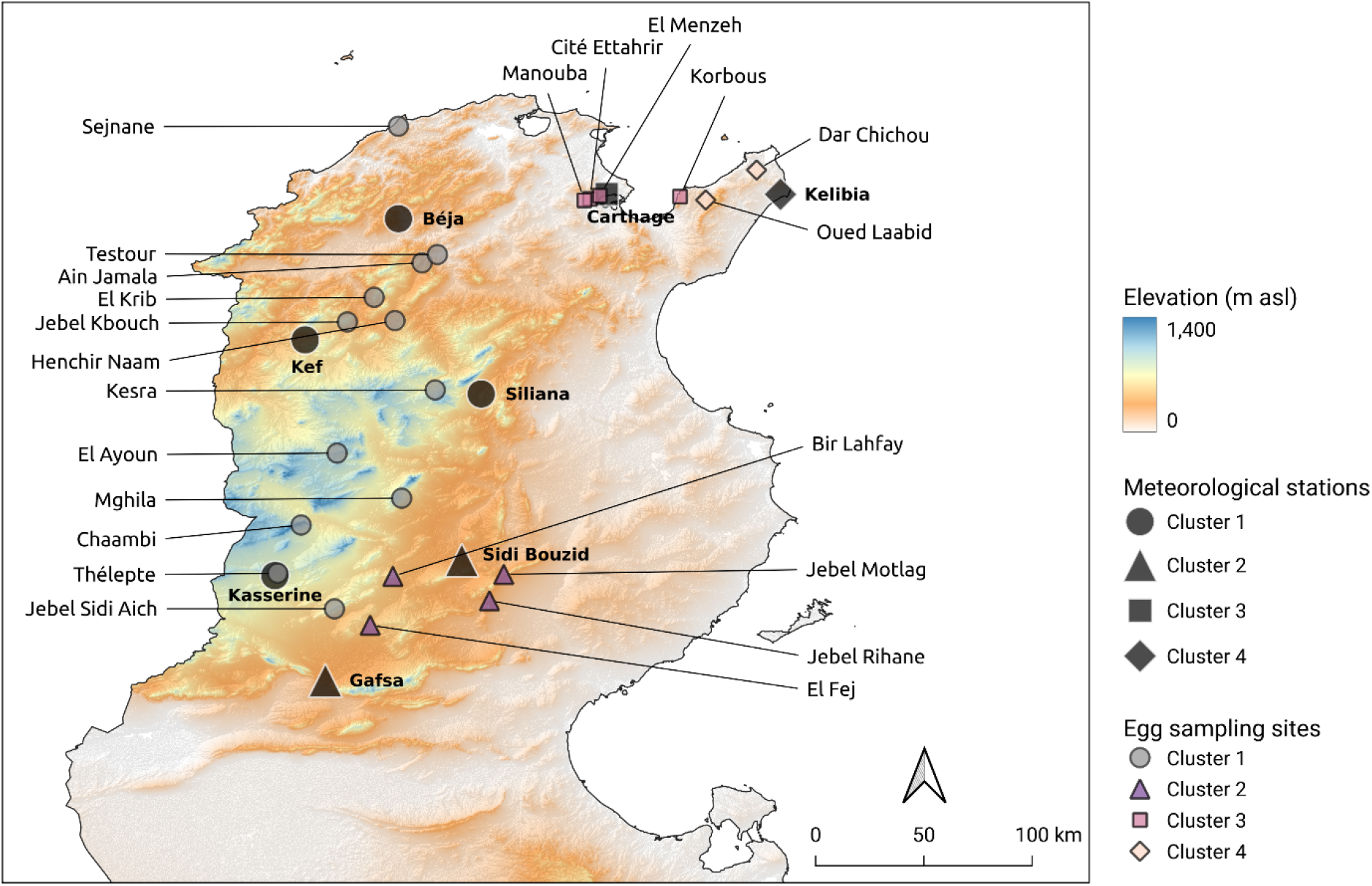
Location of egg sampling sites and meteorological stations, with associated calculated climate cluster.

**Table 1.** Collection of PPM egg masses in Tunisia (see Table SM1 for coordinates of the sites). Calculated climate clusters are indicated to represent the amount of data available per cluster.

| Site | Year of collection | Number of egg masses | Cluster | Nearest meteorological station (within 100 km and 350 m in elevation); distance |
| --- | --- | --- | --- | --- |
| Sejnane | 1995 | 20 | 1 | Béja; 51.0 km |
| Testour | 2014 | 12 | 1 | Béja; 26.3 km |
| Ain Jamala | 2010 | 15 | 1 | Béja; 26.5 km |
| El Krib | 2010 | 18 | 1 | Kef; 38.7 km |
| Henchir Naam | 1992 | 53 | 1 | Kef; 41.5 km |
| Jebel Kbouch | 1993 | 56 | 1 | Kef; 21.3 km |
| El Ayoun | 1993 | 30 | 1 | Kasserine; 73.2 km |
| Chaambi | 1995 | 27 | 1 | Kasserine; 30.3 km |
|  | 2014 | 11 |  |  |
| Thélepte | 2017 | 43 | 1 | Kasserine; 1.9 km |
|  | 2019 | 30 |  |  |
| Jebel Sidi Aich | 2014 | 31 | 1 | Kasserine; 32.7 km |
| Kesra | 2010 | 18 | 1 | Siliana; 20.8 km |
| Mghila | 2014 | 51 | 1 | Kasserine ; 71.6 km |
| Bir Lahfay | 2014 | 12 | 2 | Sidi Bouzid; 32.5 km |
| Jebel Motlag | 2017 | 38 | 2 | Sidi Bouzid; 20.6 km |
|  | 2018 | 29 |  |  |
| Jebel Rihane | 2017 | 25 | 2 | Sidi Bouzid; 25.7 km |
| El Fej | 2017 | 18 | 2 | Gafsa; 36.6 km |
| El Menzeh | 1992 | 19 | 3 | Carthage; 3.2 km |
|  | 1993 | 57 |  |  |
|  | 2014 | 10 |  |  |
| Cité Ettahrir | 2014 | 21 | 3 | Carthage; 7.9 km |
| Manouba | 2010 | 15 | 3 | Carthage; 10.5 km |
| Korbous | 1992 | 30 | 3 | Carthage; 32.5 km |
|  | 2010 | 15 |  |  |
| Dar Chichou | 1995 | 20 | 4 | Kelibia; 17.1 km |
| Oued Laabid | 1995 | 31 | 4 | Kelibia; 33.3 km |

### 2 Climate data

We used series of daily temperatures recorded (by the Institut National de Météorologie, INM, Tunis, Tunisia) in eight meteorological stations distributed within the PPM range in Tunisia (Fig. 1; Table SM2). To fill missing data in INM time series, satellite measurement of daily temperatures were also retrieved from the NASA Prediction of Worldwide Energy Resources website (https://power.larc.nasa.gov/data-access-viewer/) on the grid cells of 0.5 degree × 0.625 degree (~ 50 km × 60 km) matching the location of INM weather stations (Table SM2). The similarity of both sources of data was evaluated using Pearson correlations tests for daily maximal and daily minimal temperatures in Tunis, where the data series from INM since 1990 was the most comprehensive. Daily maximal and minimal temperatures from both data sources were found to be strongly correlated (Pearson tests, r= 0.95, *p* < 0.001 and r = 0.94, *p* < 0.001, respectively). The two types of datasets where therefore combined in case of missing data in other INM series to reconstruct uninterrupted series for the period 1990-2019 (Table SM2).

Each site of egg sampling was assigned to the nearest meteorological station (< 100 km in all cases) among those situated at an elevation within 350 meters of the egg site, an arbitrary threshold we chose to mitigate potential climatic differences along elevation gradients (Table 1, Figure SM3).

To better understand climatic features in each of the eight meteorological series (Table SM2), (i) the normal daily temperatures with seasonal contrasts over the period, as well as (ii) the overall trend since 1990, were calculated. For (i), we averaged 30 years of daily maxima (TX) and minima (TN) by day of the year, and calculated the likelihood for each day of temperatures below 0 or above 32 and 40°C, which have been suggested by Démolin (1969) and Huchon & Démolin (1970) as pivotal thresholds for phenological strategies and survival in the PPM (see also discussion in Robinet et al. 2015). For (ii), daily TX and TN were averaged per year and represented along the 30 years of data, together with the total number of days below 0 or above 32 and 40°C. Those per-station climate summaries are provided in SM4.

### 3 Statistical analyses

#### Climate clusters

The unbalanced egg sampling design throughout historical data and recent collections prevents allochronic comparisons of egg phenotypes within individual sampling sites. Therefore, we investigated climatic similarities and dissimilarities among meteorological series in order to identify regional climate clusters within which multiple meteorological series and associated egg sampling sites could be statistically grouped together. Climate-based grouping appeared more relevant and less arbitrary than using administrative regions because of the heterogeneous landscape and overall size of some regions. To do so, the monthly averages of TN and TX were calculated in each meteorological series over the period 1990-2019, resulting in a set of 24 variables (2 × 12 months) and 30 values per series (30 years). A Principal Component Analysis (PCA) was used on the covariance matrix of those variables to project the 30 years of data from each of the eight meteorological locations and better visualize their intra- and inter-group variance on reduced dimensionality. The resulting multivariate object then fed a K-medoid clustering analysis using the PAM method (Partitioning Around Medoids, see Reynolds et al. 2006, Schubert and Rousseeuw 2019) to identify relevant climate clusters (listed in Table1). The PCA could be performed on unscaled temperature variables since they were all measured in the same unit (covariance PCA), thereby giving most weight to summer months and to TX, due to higher temperature values, without neglecting other months and TN in the overall variance structure. As a consequence, the climate clusters identified using all four seasons are mostly influenced by the season eggs are exposed to (roughly June to September). Details of cluster assignation to individual points in each meteorological series are detailed in SM5. Monthly means of TN and TX of the medoid of each cluster, *i.e*., the individual point that best represents its cluster due to low average dissimilarity to all other points, are represented in SM6.

#### Interannual fluctuation of maximal summer temperature within clusters

To explore regional warming trends to which eggs are subjected within clusters over 1990-2019, the monthly means of daily maximal temperatures from meteorological series within each cluster were calculated from June to September. A linear model was then built for each cluster and each month to plot regressions over time and determine the slope for each cluster. The adequacy of residuals to Normality was checked using QQ plots.

#### Egg phenotype comparisons

Egg phenotypes could not be compared allochronically in all clusters identified because the data set was unbalanced, with only two of four clusters grouping egg samples from both periods. Further analyses on eggs are therefore focused on those two clusters, but a complementary synchronic analysis is provided in SM7 to compare egg phenotypes across all clusters within the period(s) they have in common. Since egg phenotype variables did not meet assumptions of homoscedasticity and normality for parametric tests, we used the non-parametric RANCOVA proposed by Quade (1967) to compare eggs sampled between clusters (1 and 3) and periods (1990s and 2010s). First, the response variables (Fecundity, Hatching, Sterility, Abortion and Parasitism rates; Clutch length was discarded due to its high correlation and redundancy with Fecundity) and the covariate (monthly means of TX averaged from June to September per year per cluster) were ranked separately. Second, residuals from the respective linear regression of each ranked response variable on the ranked covariate were calculated. Third, the effects of grouping factor(s) on residuals were investigated for each response variable using Quade’s RANCOVA (factors Cluster, Period, and their interaction). It was followed by pairwise t-tests and a Bonferroni correction when a significant interaction term was found.

## Results

### 1 Climate clusters

The first plane (PC1 × PC2) of the PCA performed on climatic data from all eight meteorological series based on monthly averages of TN and TX each year (n = 8 × 30 = 240 data points) accounted for 71.28% of the total inertia (Fig. 2). PAM clustering on the PCA scores indicated four relevant groups with little overlapping (Fig. 2). Cluster 1 grouped Kef, Kasserine, Siliana and Béja together, cluster 2 grouped Sidi Bouzid and Gafsa together, while cluster 3 and cluster 4 corresponded to single meteorological series, Carthage and Kélibia, respectively. Depending on the meteorological series, between 76.67 and 100% of data points (years) were correctly assigned to their cluster (SM5).

**Figure 2.**
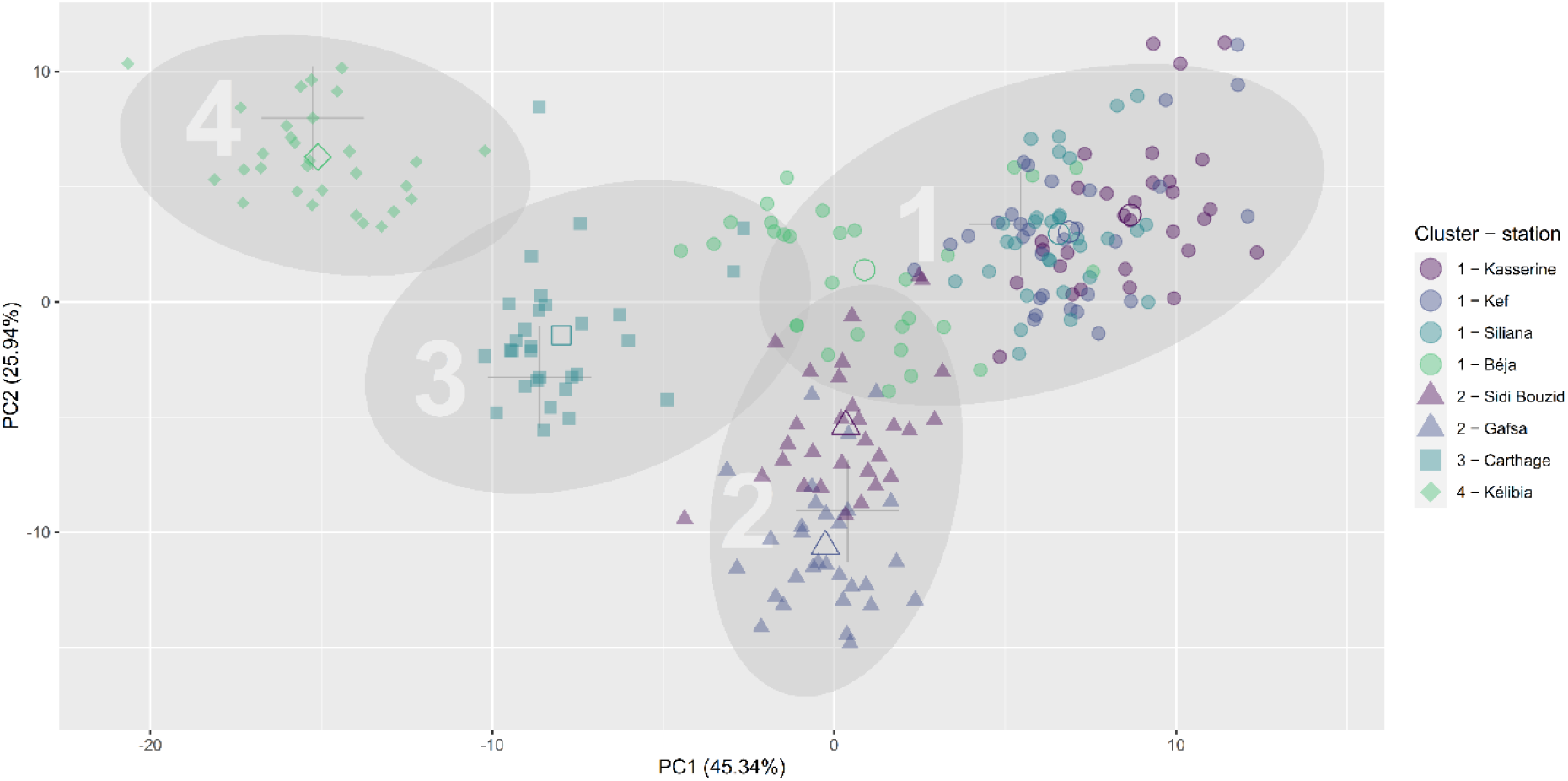
PCA scores for each year of data from the eight meteorological series (INM and NASA series, see table SM2) and 24 TN monthly average and TX monthly average variables. Each of the eight series are identified with different symbol and colour combinations. PAM clustering results are overlaid on the PCA scores with 95% confidence ellipses and different symbols for different clusters. Open points correspond to the centroids of each meteorological series, while large thin crosses mark the medoid point of each cluster.

Per-cluster climate reconstructions averaged from daily means of TX over years (Fig. 3) indicated comparatively cold winters and hot summers with a high interseasonal variability in cluster 1, warmer winters and summers in cluster 2 with similar interseasonal variability, no extreme winters or summers and lower interseasonal variability in cluster 3, and the lowest interseasonal variability with comparatively mild summers in cluster 4. Within-month variability also appeared to be the highest over the last 30 years in clusters 1 and 2. The probability to overreach 40°C in summer was found to be the highest in cluster 2, while cluster 4 showed the lowest probability of overreaching 32°C, with clusters 1 and 3 sitting in between those extremes. July and August are the warmest months in all clusters (Fig 3, Fig 4). TN monthly average within each medoid appeared to roughly reflect TX monthly average across each months of the year (SM6), indicating that similar trends can be inferred for per-cluster TN monthly average.

**Figure 3.**
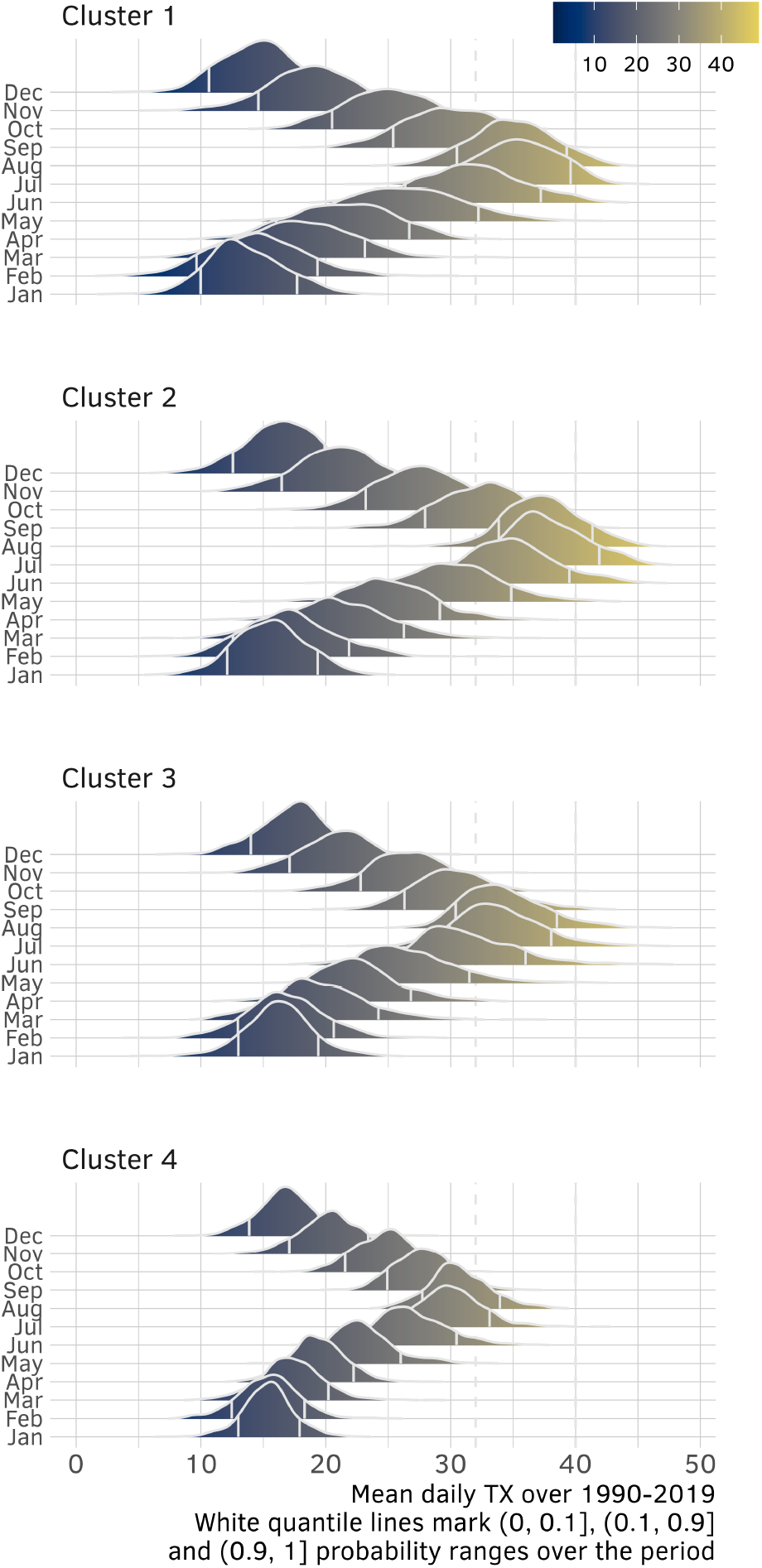
Ridge lines showing the distribution (kernel density) of daily means of TX over 1990-2019 for each cluster. The area of each distribution equals 1 and white vertical lines mark 0.1 and 0.9 quantiles.

**Figure 4.**
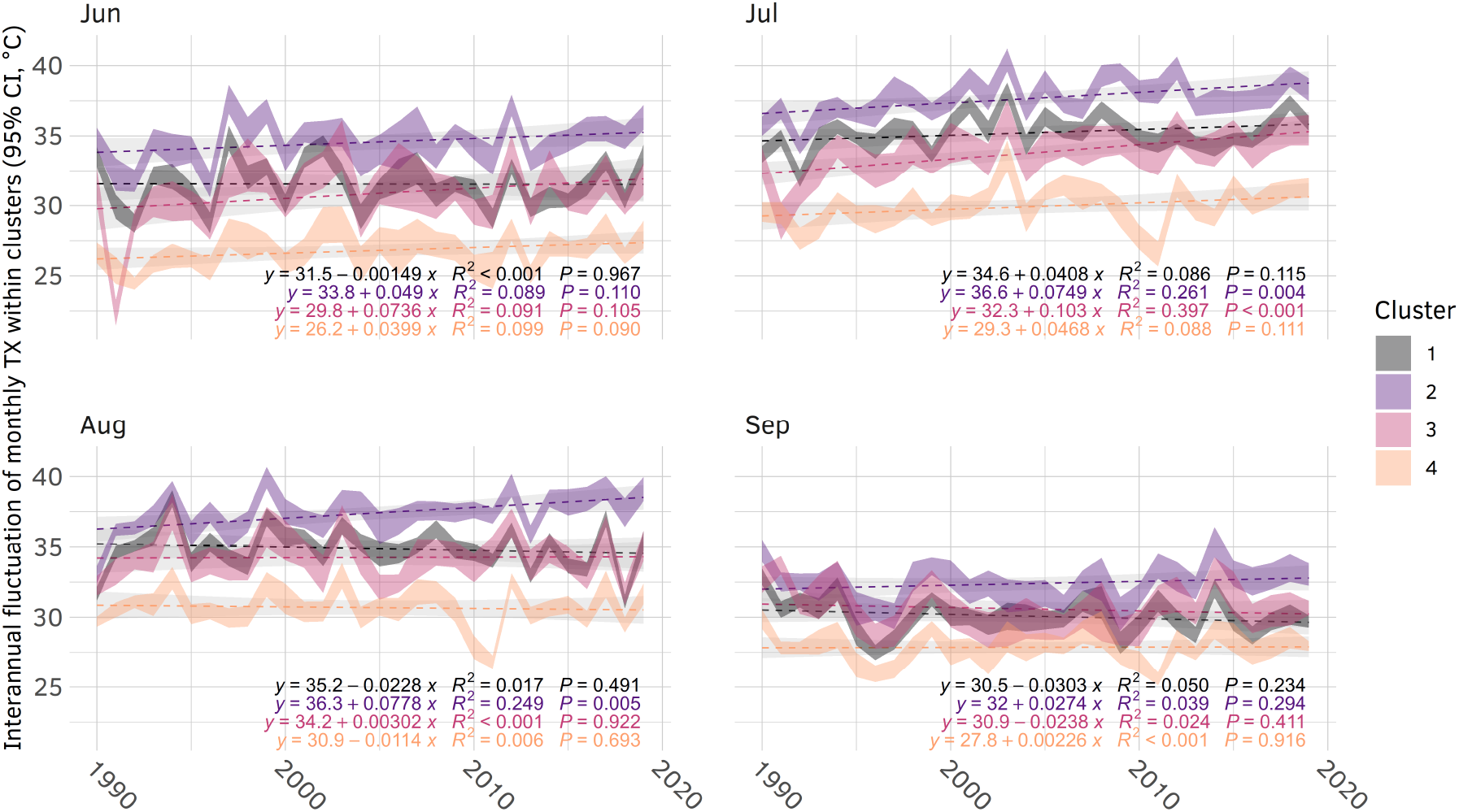
Fluctuations over time of monthly maximal temperature within each cluster, from June to September. The upper and lower lines of each ribbon correspond to 95% confidence intervals.

Fluctuations of monthly averaged TX during summer over the last three decades (Fig. 4) in each cluster indicated that September has not warmed significantly over time (no apparent slopes, non-significant linear regressions). Contrariwise, July got significantly warmer over time, particularly in cluster 2 and 3 (positive slopes, significant linear regressions). Cluster 2 appeared as the cluster with the most striking summer warming, with positive increases of TX monthly average spanning not only July, but also June (non-significant trend (p = 0.11), similar to cluster 3) and August (significant increase). Contrariwise, summers in clusters 1 and 4 appeared as the least subject to warming since 1990.

### 2 Egg comparisons

#### Clutch size

The length of egg masses was measured in case the distance between eggs of a clutch would differ among areas or periods, but this variable was found to be highly positively correlated to fecundity (Spearman correlation test, rs = 0.72, *p* < 0.001), thereby leaving little room for variations in the fecundity/length ratio. Therefore, we focused further analyses on fecundity only, which ranged from 121 ± 8.2 SE to 174.6 ± 5.9 SE among clusters and periods (Figure 5). The Quade’s RANCOVA conducted on clusters where both past and present samples have been collected (clusters 1 and 3) showed no significant difference in fecundity between those two clusters (Quade’s RANCOVA, F_df_ = 2.88_1_, p = 0.09) or periods (F_df_ = 3.01_1_, p = 0.083). However, a significant crossover interaction suggested non-parallel trends over time between them (Quade’s RANCOVA, F_df_ = 6.42_1_, p = 0.012). Indeed, cluster 3 was the only cluster where fecundity changed over time (pairwise t-test, t_df_ = 3.1_1_, p = 0.002 adjusted by Bonferroni correction), with an average decrease of 16 %.

**Figure 5.**
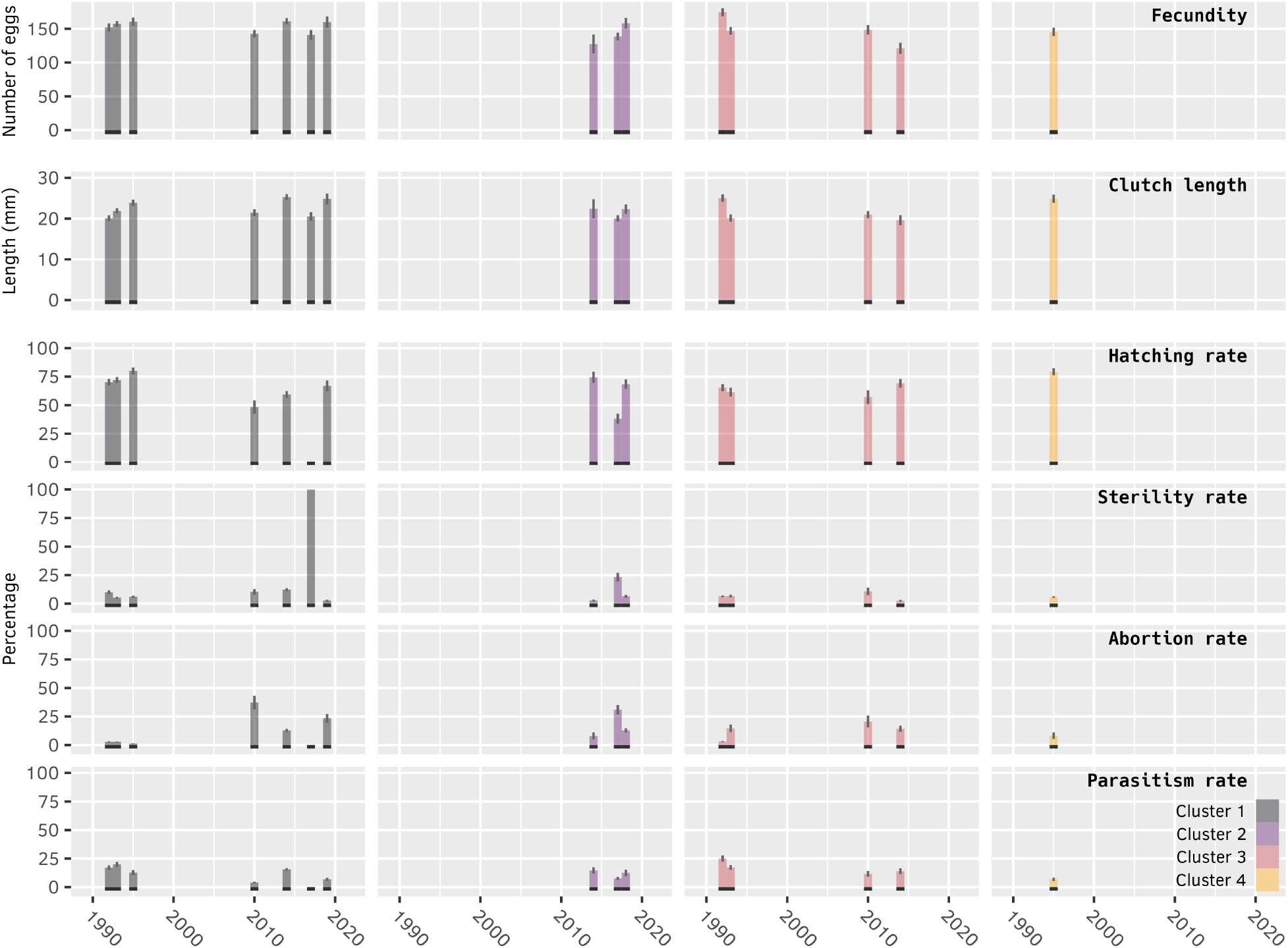
Fecundity (raw count of eggs), clutch length, hatching rate, sterility rate, abortion rate and parasitism rate per cluster and sampling year. Black markers at the bottom show years when samples have been collected, to distinguish true zeros from missing values. Error bars: SEM.

#### Hatching rate and egg mortality factors

Hatching rate did not differ significantly between clusters 1 and 3 (Quade’s RANCOVA, F_dt_ = 0.35_1_, p = 0.566), yet it significantly differed between periods (F_dt_ = 35.65_1_, p < 0.001). As found for fecundity, the interaction term was significant (F_dt_ = 20.48_1_, p < 0.001). Cluster 1 was the only cluster of the two where hatching rate decreased significantly between the past and recent periods (pairwise t-test, t_df_ = 7.5_1_, p < 0.001 adjusted by Bonferroni correction). Particularly, in 2017, hatching rate dropped to zero in cluster 1 with all eggs visually scored as sterile. Consequently, abortion and parasitism rates were also the lowest (0 %) in this cluster that year since those mortality factors can only occur at a later development stage (Figure 5, Table SM8). This average decrease over time can be attributed not only to the striking drop in 2017, but also to decreases of lower magnitude in other recent years (Figure 5). Those changes correlate negatively with the aforementioned trends in abortion rate and sterility rate. Of the two clusters, the increase in abortion rate was most prominent in cluster 1 (Figure 5), although it may be underestimated in recent years due to its null value in 2017 when eggs did not develop enough to score abortion rate.

The Quade’s RANCOVA showed that base abortion and parasitism rates differed significantly between clusters 1 and 3 (Quade’s RANCOVA, abortion rate: F_df_ = 17.13_1_, p < 0.001; parasitism rate: F_df_ = 4.53_1_, p = 0.034) and periods (abortion rate: F_df_ = 50.92_1_, p <0.001; parasitism rate: F_df_ = 20.36_1_, p < 0.001). However, the temporal increase in abortion and decrease in parasitism were similar in the two clusters since no significant interaction was observed (abortion rate: F_df_ = 0.08_1_, p = 0.776; parasitism rate: F_df_ = 1.34_1_, p = 0.248)..

Finally, to evaluate the influence of the 2017 peculiarity on overall trends in cluster 1 and investigate other changes that may have been concealed by this heatwave, another Quade’s RANCOVA has been performed on a subset of the data without cluster 1 in 2017 (see descriptive statistics in Table SM9). It revealed that hatching and abortion rates still changed over time in the two clusters, to a lower extent than when considering data from 2017 in cluster 1 (hatching rate: F_df_ = 8.2_1_, p = 0.004; abortion rate: F_df_ = 7.6_1_, p = 0.006), whereas sterility and parasitism did not change in any of the two clusters (sterility: F_df_ = 1_1_, p = 0.315; parasitism: F_df_ = 0.3_1_, p = 0.563).

## Discussion

Climate change has been recognized to be one of the major phenomena that may affect forest insect populations (Jactel et al., 2019; Ramsfield et al., 2016). While many studies reported the occurrence of more frequent and larger insect outbreaks (Raffa et al., 2008; Robinet & Roques, 2010), the opposite have also been observed (Pureswaran et al., 2018; Rozenberg et al., 2020). Ongoing global warming may exert mixed effects on population dynamics (Dreyer & Baumgärtner, 1996; Huang et al., 2008), and ultimately have an impact on species distributions, as has been observed with the PPM northern range expansion in Europe (Battisti et al. 2005) and the southern range retraction in Tunisia (Bourougaaoui et al. 2021). A report by the German Technical Cooperation Agency (GTZ et al. (2007)) has predicted more intense and longer heatwaves in Tunisia, with temperatures tending to rise even further in the coming century. To better understand potential adverse effects of climate change at the southern edge of the PPM range, the present study sought to explore variations in egg survival and hatching and their potential relationship with climate variations, based on a set of historical and recent field samplings across Tunisia.

### i. Hatching failure and heatwaves

The decrease in hatching rate observed in the 2010s period was caused by a steep increase in the rate of sterile eggs and a clear increase in abortion rate (i.e., fertile eggs with failed embryo development), the latter being possibly related to warmer conditions during embryonic development. The strikingly high sterility rate observed in 2017 could be related to an unusually long series of 10 consecutive days above 40 °C recorded that year (see Fig. 6 and Fig. SM10 for meteorological data from the closest station of the site sampled that year). While extreme compared to the last 30 years, this anomaly reflects the global increase in the total number of acute heat days recorded in August in this station (Fig. 6). This overall trend in turn corroborates the assumption that the likelihood of such stochastic events should increase with future climate change and cannot be neglected since they might represent a prime cause of mortality in the PPM, before the average warming. We found that July is the most rapidly warming month in Tunisia, but egg masses in sites within cluster 1 are mostly laid after July and occur in August. Since all 43 egg masses from cluster 1 in 2017 were collected in late August, after this long heatwave, egg development may have been directly impacted before any sign of embryogenesis could be detected (noted as “sterile” from visual inspections). Such acute heat may also have accelerated pheromone decay due to higher evaporation rate, hence affecting mating success and egg fecundation in the first place (Groot & Zizzari, 2019; Linn et al., 1988), or adult gametes (Sales et al., 2018). These results bear a close resemblance to those shown by Rocha et al. (2017), which revealed that negative effects appeared on Tunisian egg masses at 42°C after only 3 days of heatwaves, and no survivorship was noted at 44°C.

**Figure 6.**
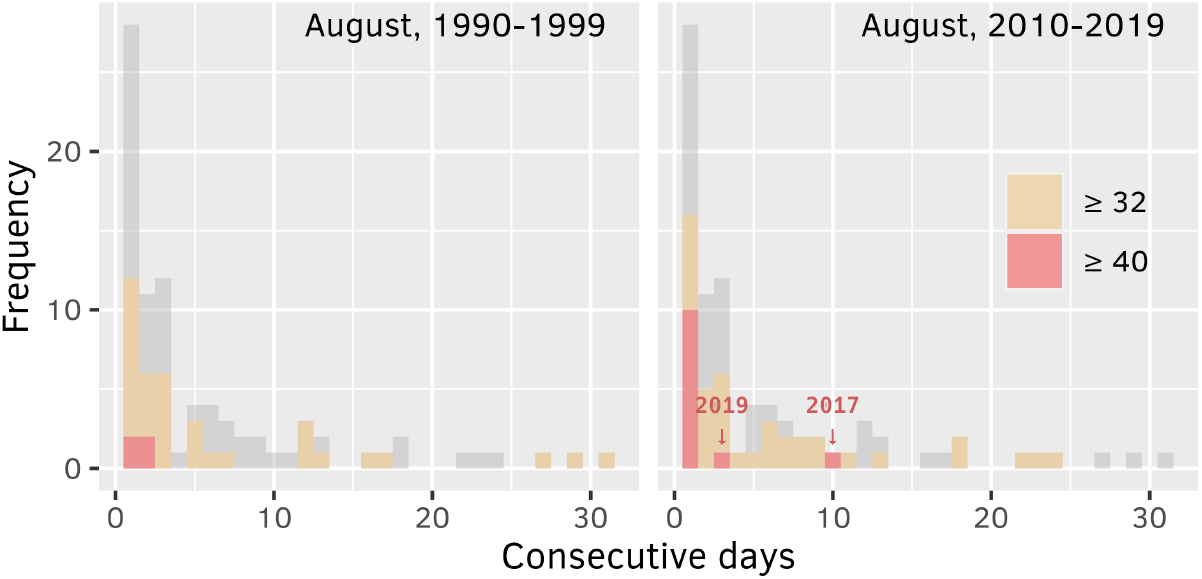
Frequency of consecutive days in August overreaching 32 (orange) and 40°C (red) over 1990-1999 and 2010-2019 in the Kasserine station. This station is situated less than two kilometers away from Thélepte, the egg sampling site where 43 egg clutches (100 %) were sterile in 2017. Grey histograms correspond to the total distribution across both periods.

Hatching rate was slightly higher in cluster 2 than cluster 1 in the 2010s (Table SM8), which can be attributed to the drop in 2017 in cluster 1 (Figure 5). Despite being true for all egg masses in cluster 1 that year, caution must be taken before generalizing the low egg survival recorded in 2017 to a temporal trend in the whole cluster 1. First, all eggs of cluster 1 in 2017 originated from the single site of Thélepte, however egg masses were collected on multiple trees scattered across the area, thereby reducing the risk of biased sampling, and the sample size was one of the largest of the whole data series (43 eggs masses; see Table 1). Second, inter-annual variability in climate and other factors not measured in this study cannot be neglected, meaning that the recent years cannot be summarized to the year of 2017 which has been shown to be extreme. However, global climatic models as well as the data presented in this manuscript suggest that these types of events are likely to increase in frequency with climate change, together with average warming, and therefore should be explicitly taken into account when analyzing PPM dynamics because they represent prime causes of lethality. By putting together long data series, the present study provides a broader view on the spatial and temporal variations in the fecundity penalty that may result from these extremely high temperatures, as well as how the timing of these heatwaves may differentially impact populations from different areas. Beyond the dramatic drop in egg survival observed in 2017 in cluster 1, smaller penalties on hatching rate have also been observed during other years of the 2010s. Those smaller decreases of egg survival may have a snowball effect on whole colony success, as shown in Spain by Pérez-Contreras et al. (2003) who found that 32 individuals is a threshold above which larval growth reaches its maximum and mortality drops substantially. A similar result was observed in an exploratory experiment in a French population during winter, where survival was null for colonies of less than 50 individuals (Roques et al., 2015).

The probability to overreach biological thresholds was found to be the highest in cluster 2 and to significantly increase over time, but no historical egg data from this cluster are available to confirm whether egg survival used to be higher in the 1990s. Our finding showing that August conditions in 2017 have likely been lethal in Thélepte (and possibly to a largest extent in cluster 1) may indicate that similar or worse dramatic effects associated with stochastic meteorological events are to be expected in cluster 2 where summers are already both the hottest and the most rapidly warming (Figures SM4). However, despite a longer and more intense 2017 heatwave in cluster 2 (Figure SM11), eggs occur later (hatching starting in mid-October for recent samples) than in cluster 1 (hatching starting in early September), which may mitigate summer heatwave threats thanks to phenological avoidance. This contrasting situation warrants the high relevance of future research in populations from cluster 1 and 2 to observe in real-time and test how climate change may cause the retraction of the PPM at its southern edge (see Bourougaaoui et al. 2021).

Temperature thresholds of 32°C and 40°C have been frequently used in the literature as pivotal for range limits of the PPM. They have been inferred from observations of survival in nature, but also appear consistent with more recent experiments in multiple populations of the PPM. Eggs from a French population were able to withstand a short transient exposure to a daily maximal temperature of 40°C during several consecutive days with no mortality impact (Robinet et al., 2013), while eggs from a nearby population could survive a single 6-hour-long exposure to up to 44°C (Poitou, 2021). However, mortality appeared on egg masses from a Tunisian population at 42°C after only three days of 4-hour daily exposures (Rocha et al., 2017). While slightly different methods have been used and make it difficult to compare populations, those results help narrowing down the tipping point at which PPM egg survival is impacted, depending on the duration of exposure. Regarding larvae, the survivorship of L1 and L2 from Portuguese populations started to drop after 4-hour exposures to 36°C and 40°C, respectively (Santos et al., 2011), showing higher susceptibility in early larvae compared to eggs. Recently, Poitou et al. (2022) determined experimentally the thermal performance curves in development rate in the first four larval instars in a French population, highlighting that the 32°C threshold is above the optimal development temperature, within the decreasing performance phase. The 32°C and 40°C thresholds proposed by Démolin (1969) and (Huchon & Démolin, 1970) thus appear as conservative but consistent integrators of whether a population is facing stressful conditions regardless of the exact duration of exposure, which our results corroborate with full mortality after the 2017 heatwave but not after the intense yet less stringent 2019 heatwave (Figure SM10).

In many parts of the world, climate warming more readily impacts nighttime than daytime, thereby contributing to a decrease in the diurnal thermal range (DTR; Higashi et al., 2020; see also *e.g*., Béja, Kef and Carthage in SM4). Several studies (*e.g*., Higashi et al., 2020; Zhao et al., 2014) have found that the impact of heat stress endured throughout the day on fitness can be exacerbated by increasing nighttime temperatures, since thermal fluctuations can help ectotherms repairing or buffering thermal injuries. In the pine processionary moth, however, warmer winters have facilitated larval feeding activity in France, which occurs when daytime and nighttime temperatures exceed 9°C and 0°C, respectively (Battisti et al., 2005). Likewise, the unusually warm night temperatures during the 2003 heatwave in southern Europe benefited to the PPM in the Alps by eliciting dispersal of imagos, which are nocturnal (Battisti et al., 2006). This may be attributed to lower heat intensity due to altitude, and delayed phenology in the Alps which made adults the exposed stage. Due to their nocturnal activity and short lifespan, the beneficial influence of TN on flight activity may have prevailed in adults, whereas eggs or young larvae in other areas with a more advanced phenology are more likely to suffer from prolonged exposure to both high TX (causing thermal stress) and high TN (limiting recovery from heat injury; Zhao et al., 2014). Little is known about the impact of warming TN in Tunisia where the average temperature is higher, and warming nights may impede the capacity to recover from heat stress, but these findings suggest that local phenologies are key to determining the impact of reduced diurnal thermal fluctuations in the PPM.

### ii. Thermal tolerance and phenology among populations

Temperature is a crucial abiotic factor that can lead to local adaptations in insects and shape their geographic range (Bush et al., 2016; Hoffmann et al., 2002; Kellermann et al., 2012; Sinclair et al., 2012). Various thermal environments may therefore be associated with differences in behaviours and even physiological tolerance (Calosi et al., 2010). Numerous studies have focused on geographical variation in thermal tolerance in the fruit fly *Drosophila melanogaster*, considered as one of the most widely distributed insect species, highlighting local variation in the thermal tolerance and performance of distinct populations (see e.g., Sinclair et al., 2012). In this species, Hoffmann et al. (2002) found opposing clines in resistance to temperature extremes when comparing numerous Australian populations along a gradient from tropical to temperate latitudes, which suggests that thermal tolerance traits are under direct climatic selection. Likewise, summer temperature has been hypothesized as being a strong selection pressure in the PPM due to the mortality observed in eggs (Rocha et al. 2017) and neonate larvae (Santos et al. 2011) after experimental heat exposure, combined with the wide range of climates under which the PPM occurs. Variations in environmental parameters may induce phenological shifts in all stages of this species either as a consequence of altered developmental time or responses to stressors (Berardi et al., 2015; Robinet et al., 2015). In areas where summers are the warmest, early mortality caused by heat stress may favour late-emerging individuals through natural selection, especially as adult females are shortlived and lay eggs only once (Rocha et al. 2017). A striking example of phenological differentiation among PPM populations was recently discovered in Portugal, where two sympatric forms exhibit contrasted life cycles: one with the typical overwinter larval development, and one with a summer larval development (Santos et al., 2011). While little is known on the causal factors that promoted the emergence of both forms in the same area, Godefroid et al. (2016) demonstrated that the range of the summer population is restricted to central-coastal Portugal due to climatic constraints, while winter populations expands northward. This may be due to the comparatively low thermal resistance found in eggs of the summer form, which develop into mature larvae before the peak of summer, as opposed to eggs and early instar larvae of the winter-developing form which usually occur in the middle of summer and are more likely to be exposed to heats (Rocha et al. 2017). The increase in climatic variability is expected to exert effects on insect species that differ from those caused by gradual global warming (Schreven et al., 2017). Large-scale heatwaves are known to have contrasted effects on different populations of the PPM depending on local climates and phenologies (Robinet et al., 2015). In Europe, the summer heatwave in 2003 led to a collapse of PPM populations in northern areas in France (Robinet et al., 2013), while it benefited to adult dispersal and altitudinal expansion in the Italian Alps, likely due to flight thermal thresholds more easily met than usual near the elevation edge (Battisti et al., 2006). These findings are congruent with the available data in Tunisia where phenology in cluster 2 is delayed compared to that in cluster 1, most likely due to the climatic and altitudinal differences found between those clusters. This fact could contribute to explain the lower hatching rate found in cluster 1 than in cluster 2 in 2017 (Figure 5), since eggs and neonate larvae occur after most summer heats in cluster 2 as a result of later adult emergences. This suggests that populations from the warmest areas of Tunisia may not be the most vulnerable to climate change thanks to phenological adaptation favouring heat avoidance, as opposed to populations from areas where individuals are close to their physiological limits but did not evolve differentiated phenology in response to heat lethality.

Despite the impacts of the PPM on Aleppo pine forests in Tunisia, little is known about how climate change can alter the phenology of Tunisian population and whether it is is spatially structured by climate heterogeneity. According to Robinet et al. (2015), predictable unfavourable conditions to which PPM populations are exposed can be alleviated by differentiated phenologies (phenological plasticity and/or adaptations), which is supported by later adult emergences in cluster 2 on average. By contrast, unpredictable adverse conditions may be mitigated by bet-hedging strategies such as prolonged diapause, as it creates heterogeneity within populations and siblings by expanding the life cycle of only a fraction of individuals that stay sheltered in the ground at the pupal stage and evade climate stochasticity (Le Lann et al., 2021; Salman et al., 2019). Diapausing individuals may therefore contribute to reconstituting local population collapses after extreme events, but the cost is that survival decreases with the total duration of diapause due to other factors of mortality (fungi, pupal predation, etc.) (Salman et al. 2019). A significant relationship was found between the rate of prolonged diapause and both cold and warm winter temperatures, presumably because they serve as cues of the likelihood of either cold or hot lethal temperatures caused by interannual climate uncertainty (Salman et al. 2019). The acute heat stress exerted on PPM egg masses in Tunisia may help explain the retraction of this pest from southernmost regions, and further investigations on phenological differentiations and prolonged diapause strategies among populations incurring different levels climate harshness are now necessary to predict further distribution changes.

### iii. Host-parasitoid interactions and outbreaks in a warming climate

A growing body of literature reveals that parasitoids are often more sensitive to climate warming than other trophic levels because of their higher position in the food web (Jeffs & Lewis, 2013; Rosenblatt & Schmitz, 2016). Climate change can lead to phenological asynchrony between parasitoids and their hosts in cases where the phenology of the interacting species respond variously to the same climatic cue (Visser & Holleman, 2001), or when the interacting species use different cues to initiate emergence or development (Jeffs & Lewis, 2013; Walther, 2010). Phenological asynchrony may also appear if one of the interacting partners rapidly develops or has a seasonal diapause in response to warming (Forrest, 2016). Parasitism rates therefore tend to decrease with increasing climatic variability that impedes parasitoids from tracking host populations (Chidawanyika et al., 2019). Alternatively, but not exclusively, eggs may escape parasitic attacks thanks to various counter-adaptations whose relative weight may differ among regions and populations. The identification of eggs by natural enemies may be hindered making egg masses inconspicuous, as PPM female covering eggs by greyish-brown scales similar to the colour of pine twigs (Battisti et al., 2015). These scales function not only as visual protection but also as factors impacting the microclimate of eggs and therefore their development rate (Milani, 1990), and as physical barriers limiting parasitoids from locating individual eggs in the clutch and greatly reducing their success (Pérez-Contreras & Soler, 2004). The chief defence against parasitoids, widely discussed in insects and in particular *Drosophila spp*., is encapsulation followed by melanisation, an immune responses which sequesters and kills foreign body (Cavigliasso et al., 2021; Wertheim et al., 2005). Such immune system with specific cells (hemocytes) is well-developed in larval stages as already observed in PPM larvae (Boudjahem et al., 2019), however, a study conducted by Reed et al. (2007) showed that hosts in the egg stage can mount a cellular immune response against parasitoid eggs and larvae (Reed et al., 2007). Research on a wide range of species reveals that small changes in temperature can significantly shape insect immunity as well as parasitoid fitness (Murdock et al., 2012). Increases in temperature can consequently promote or repress the encapsulation process, which in turn may influence the outcome of parasitic success, (Cavigliasso et al., 2021). The outbreaks of phytophagous insects are expected to increase in the future as parasitism decreases (Stireman et al., 2005). At the southern edge of PPM distribution (North Africa), some studies showed that extremely high temperatures could both disrupt population regulation mechanisms and decrease the severity of outbreaks (Bouzar.Essaidi et al., 2021; Pureswaran et al., 2018). The lower PPM fecundity in this area (when compared to that at the northern edge, in southern Europe) due to lower plant quality (Bouzar.Essaidi et al., 2021; Pimentel et al., 2010) may explain why egg parasitism is also lower with climate warming. Our results showed that the mean number of eggs per egg mass (155 ± 2.1 in cluster 1, 142 ± 4.5 in cluster 2, 150 ± 3.5 in cluster 3, 145 ± 6.3 in cluster 4; 151 ± 1.7 when merging all clusters) was considerably lower than that found in Bulgaria (226 ± 43.2) and France (194.3 ± 50.1) (Georgiev et al., 2020). The average rate of parasitism also showed a different pattern than what was found in northern parts of PPM distribution, with increases in coastal regions namely in cluster 1 in Tunisia, while it decreases in France and Spain from core to front populations and along an altitudinal gradient, respectively (Georgiev et al., 2020; Hódar et al., 2021). Although parasitoids account in egg mortality, their influence here was low compared to other factors (sterility and abortion) (Figure 5), and is therefore unlikely to be the main driver of PPM collapses at the southern edge of the distribution. Significant local warming in Tunisia appears as a prime candidate factor contributing to the sharp decrease of PPM populations (Bourougaaoui et al., 2021).

### iv. Other factors influencing distribution

Factors other than climate warming may putatively affect the survival and persistence of the PPM at its southern range edge. Embryonic mortality can be impacted by excessive exposure to intense solar radiation, particularly in southern parts of the distribution, as PPM females tend to lay their eggs exposed to the sun (Démolin, 1969). Another factor often modulating the spatial occurrence of insects is food availability. Nevertheless, it is rarely a limiting factor in the PPM because larvae feed on evergreen trees that are well distributed in the environment, from natural or semi-natural stands to urban areas where they often occur in relatively high numbers as ornamental trees (Martin, 2005). Natural enemies such as pathogens or predators (mostly insect parasitoids) at early larval stages have been suspected to cascade into increasing mortality during larval development because of the impact on the colony size and silk weaving effort to build a tent (Auger Rozenberg et al., 2015; Roques et al., 2015), however (1) there is no evidence that the enemy pressure would differ among areas investigated here, and (2) temperature, particularly summer heat waves or early autumnal cold snaps, are often put forward as a major cause of early mortality (Robinet et al., 2015).

### v. Conclusion

Heat tolerance has received close attention in insects, however its fluctuation throughout ontogeny and effects persisting from one developmental stage to another are still poorly documented. Besides the PPM, few case studies showed that the effects of acute heat stress received early in life cycle may be carried over to later instars. This was demonstrated in holometabolous insects such as the tropical butterfly, *Bicyclus anynana* (Klockmann et al., 2017). Beside consequences of heat on immediate mortality investigated in experimental work (e.g., Rocha et al 2017) or inferred in the present study by putting together long time series, the ultimate fitness of individuals that survive challenging heats at the egg stage or first larval instar would therefore be of great interest to understand the impacts of climate warming at the southern edge of the PPM. This insect remains one of the ideal models to study these questions owing to (i) the availability of historical data, (ii) its already demonstrated spatial and phenotypic causal response to climate change (Battisti et al., 2005; Robinet et al., 2007; Poitou et al., 2022), and (iii) ongoing processes at play in its southernmost distribution affecting population persistence (Bourougaaoui et al. 2021; this study).

## Supporting information

Supplementary material (11 items)

## Acknowledgements

We are grateful to Adel Ben Abada (INRGREF, Tunis, Tunisia) for his valuable help in the field. We acknowledge the National Institute of Meteorology (INM) in Tunis for providing temperature datasets. Version 5 of this preprint has been peer-reviewed and recommended by Peer Community In Ecology (https://doi.org/10.24072/pci.ecology.100097).

## Author contributions

Conceptualization: AB, CR, MLBJ, ML; Data curation: AB; Formal analysis: AB, ML; Funding acquisition: CR, MLBJ; Investigation: AB; Methodology: AB, CR, ML; Project administration: CR, MLBJ; Supervision: CR, MLBJ, ML; Writing – original draft: AB, CR, ML; Writing – review and editing: AB, CR, MLBJ, ML.

## Data, script and code availability

Data and R scripts are publicly available at https://doi.org/10.15454/RUEIOA.

## Supplementary information

Supplementary tables and figures are available at https://www.biorxiv.org/content/10.1101/2021.08.17.456665v5.supplementary-material:

**Table SM1.** Coordinates of sampling sites.

**Table SM2.** Temperature datasets (combination of data from the Institut National de Météorologie, INM, and the NASA data in corresponding grid cells of 0.5 degree × 0.625 degree (roughly 50 × 60 km)) and coordinates of meteorological stations.

**Figure SM3.** Geographic and elevational distances between meteorological stations and egg sampling sites (lower opacity if >100 km and >350 m, respectively).

**Figure SM4.** Climate data in eight Tunisian regions between 1990 and 2019 (data source: INM and NASA, see Table SM2). Left charts show the mean year in each region by averaging daily maxima (red) and minima (blue) by day of the year over the period, represented as 95% CI ribbons. Right charts show the yearly average of daily maxima (red) and minima (blue), represented as 95% CI ribbons, and corresponding Theil-Sen estimators. Thick grey ribbons in the background show the maximal thermal range across all nine regions depending on day of the year (left) or year (right). The bottom part of left charts shows the likelihood of temperatures ≤ 0 (blue), ≥ 32 (beige) or ≥ 40 °C (red), while the bottom part of right charts shows the annual number of days below or above those thresholds. The 366th day during leap years was discarded due to its lower sample size.

**Figure SM5.** Number of years assigned to the different clusters for each meteorological series (PAM clustering on data from January to December).

**Figure SM6.** TX and TN from January to December in each cluster medoid (PAM clustering on data from January to December).

**SM7.** Synchronic comparison of egg phenotypes among all four clusters.

**Table SM8.** Descriptive statistics with all data including cluster 1 in 2017: observed mean (M), Quade’s adjusted mean (Madj) and associated standard error (SE) for the different response variables.

**Table SM9.** Descriptive statistics without data from cluster 1 in 2017: observed mean (M), Quade’s adjusted mean (Madj) and associated standard error (SE) for the different response variables.

**Figure SM10.** August daily maximal temperature recorded in 1990-2019 at the Kasserine station, near the sampling site of Thélepte (cluster 1). Yellow points correspond to daily maxima ≥ 32 °C, red points correspond to daily maxima ≥ 40 °C. Smooth lines are fitted with the “gam” (generalized additive model) method.

**Figure SM11.** Daily maximal temperature recorded in August 2017 in all stations. Béja, Kasserine, Kef and Siliana belong to Cluster 1; Gafsa and Sidi Bouzid belong to Cluster 2; Carthage belongs to Cluster 3; Kélibia belongs to Cluster 4. Yellow points correspond to daily maxima ≥ 32 °C, red points correspond to daily maxima ≥ 40 °C. Smooth lines are fitted with the “gam” (generalized additive model) method.

## References

Allen, S., Cardona, O., Cutter, S., Dube, O. P., Ebi, K., Handmer, J., Lavell, A., Mastrandrea, M., McBean, G., Mechler, R., & Nicholls, N. (2012). Managing the Risks of Extreme Events and Disasters to Advance Climate Change Adaptation. Special Report of Working Groups I and II of the Intergovernmental Panel on Climate Change. In. https://doi.org/10.13140/2.1.3117.9529

Auger Rozenberg, M. A., Barbaro, L., Battisti, A., Blache, S., Charbonnier, Y., Denux, O., Garcia, J., Goussard, F., Imbert, C.-E., Kerdelhué, C., Roques, A., Torres Leguizamon, M., & Vetillard, F. (2015). Ecological Responses of Parasitoids, Predators and Associated Insect Communities to the Climate-Driven Expansion of the Pine Processionary Moth. In A. Roques (Ed.), Processionary Moths and Climate Change: An Update (pp. 311–357). Springer Netherlands. https://doi.org/10.1007/978-94-017-9340-7_7

Battisti, A., Avci, M., Avtzis, D. N., Jamaa, M. L. B., Berardi, L., Berretima, W., Branco, M., Chakali, G., El Alaoui El Fels, M. A., Frérot, B., Hódar, J. A., Ionescu-Mălăncuş, I., ípekdal, K., Larsson, S., Manole, T., Mendel, Z., Meurisse, N., Mirchev, P., Nemer, N., … Zamoum, M. (2015). Natural History of the Processionary Moths (Thaumetopoea spp.): New Insights in Relation to Climate Change. In A. Roques (Ed.), Processionary Moths and Climate Change: An Update (pp. 15–79). Springer Netherlands. https://doi.org/ https://doi.org/10.1007/978-94-017-9340-7_2

Battisti, A., Stastny, M., Buffo, E., & Larsson, S. (2006). A rapid altitudinal range expansion in the pine processionary moth produced by the 2003 climatic anomaly. Global Change Biology, 12(4), 662–671. https://doi.org/ https://doi.org/10.1111/j.1365-2486.2006.01124.x

Battisti, A., Stastny, M., Netherer, S., Robinet, C., Schopf, A., Roques, A., & Larsson, S. (2005). Expansion of geographic range in the pine processionary moth caused by increased winter temperatures. Ecol Appl, 15(6), 2084–2096. https://doi.org/ https://doi.org/10.1890/04-1903

Ben Jamâa, M., & Jerraya, A. (1999). Essai de lutte contre la processionnaire du pin, Thaumetopoea pityocampa Schiff.(Lep., Thaumetopoeïdae), à l’aide de Bacillus thuringiensis Kurstaki (ECOTECH-PRO). Annales de l’INRGREF,

Berardi, L., Branco, M., Paiva, M., Santos, H., & Battisti, A. (2015). Development time plasticity of the pine processionary moth (Thaumetopoea pityocampa) populations under laboratory conditions. Entomologia, 3, 19–24. https://doi.org/10.4081/entomologia.2015.273

Boudjahem, I., Brivio Fransisco, M., Berchii, S., Mastore, M., & Aoouati, A. (2019). Identification and Quantification of the Most Abondant Hemocytes in the Pine Processionary Caterpillar; ThaumetopoeaPityocampa (Notodontidae). Energy Procedia, 157, 992–998. https://doi.org/ https://doi.org/10.1016/j.egypro.2018.11.266

Bourougaaoui, A., Jamâa, M. L. B., & Robinet, C. (2021). Has North Africa turned too warm for a Mediterranean forest pest because of climate change? Climatic Change, 165(3-4), 46. https://doi.org/10.1007/s10584-021-03077-1

Bouzar.Essaidi, K., Branco, M., Battisti, A., Garcia, A., Fernandes, M. R., Chabane, Y., Bouzemarene, M., & Benfekih, L. (2021). Response of the egg parasitoids of the pine processionary moth to host density and forest cover at the southern edge of the range. Agricultural and Forest Entomology, 23(2), 212–221. https://doi.org/ https://doi.org/10.1111/afe.12423

Bush, A., Mokany, K., Catullo, R., Hoffmann, A., Kellermann, V., Sgrò, C., McEvey, S., & Ferrier, S. (2016). Incorporating evolutionary adaptation in species distribution modelling reduces projected vulnerability to climate change. Ecology Letters, 19(12), 1468–1478. https://doi.org/ https://doi.org/10.1111/ele.12696

Calosi, P., Bilton, D. T., Spicer, J. I., Votier, S. C., & Atfield, A. (2010). What determines a species’ geographical range? Thermal biology and latitudinal range size relationships in European diving beetles (Coleoptera: Dytiscidae). Journal of Animal Ecology, 79(1), 194–204. https://doi.org/10.1111/j.1365-2656.2009.01611.x

Carus, S. (2009). Effects of defoliation caused by the processionary moth on growth of Crimean pines in western Turkey. Phytoparasitica, 37(2), 105–114. https://doi.org/10.1007/s12600-008-0018-z

Cavigliasso, F., Gatti, J. L., Colinet, D., & Poirié, M. (2021). Impact of Temperature on the Immune Interaction between a Parasitoid Wasp and Drosophila Host Species. Insects, 12(7), 647. https://www.mdpi.com/2075-4450/12/7/647

Charmantier, A., & Gienapp, P. (2014). Climate change and timing of avian breeding and migration: evolutionary versus plastic changes. Evolutionary applications, 7(1), 15–28. https://doi.org/10.1111/eva.12126

Chidawanyika, F., Mudavanhu, P., & Nyamukondiwa, C. (2019). Global Climate Change as a Driver of Bottom-Up and Top-Down Factors in Agricultural Landscapes and the Fate of Host-Parasitoid Interactions [Review]. Frontiers in Ecology and Evolution, 7(80). https://doi.org/10.3389/fevo.2019.00080

Chuine, I. (2010). Why does phenology drive species distribution? Philosophical Transactions of the Royal Society B: Biological Sciences, 365(1555), 3149–3160. https://doi.org/ doi:10.1098/rstb.2010.0142

Chuine, I., de Cortazar-Atauri, I. G., Kramer, K., & Hänninen, H. (2013). Plant Development Models. In M. D. Schwartz (Ed.), Phenology: An Integrative Environmental Science (pp. 275–293). Springer Netherlands. https://doi.org/10.1007/978-94-007-6925-0_15

Clark, B. R., & Faeth, S. H. (1997). The consequences of larval aggregation in the butterfly Chlosyne lacinia. Ecological Entomology, 22(4), 408–415. https://doi.org/ https://doi.org/10.1046/j.1365-2311.1997.00091.x

Coumou, D., & Rahmstorf, S. (2012). A decade of weather extremes. Nature Climate Change, 2(7), 491–496. https://doi.org/10.1038/nclimate1452

Crozier, L. (2004). WARMER WINTERS DRIVE BUTTERFLY RANGE EXPANSION BY INCREASING SURVIVORSHIP. Ecology, 85(1), 231–241. https://doi.org/ https://doi.org/10.1890/02-0607

Démolin, G. (1965). Grégarisme et subsocialité chez Thaumetopoea pityocampa Schiff. Nids d’hiver– activité de tissage. Actes du Vº Congress de L’Union Internationale pour L’étude des insectes Sociaux,

Démolin, G. (1969a). Bioécologie de la processionnaire du pin Thaumetopoea pityocampa Schiff. Incidences des facteurs climatiques. Boletin del Servicio de Plagas Forestales (23), 9–24. https://hal.inrae.fr/hal-02732616

Démolin, G. (1969b). Comportement des adultes de Thaumetopoea pityocampa Schiff. Dispersion spatiale, importance écologique. Annales des sciences forestières,

Démolin, G., & Rive, J. (1968). La processionnaire du pin en Tunisie. Ann. I.N.R.F. Tunisie, 1(1), 1–19.

Denno, R., & Benrey, B. (1997). Aggregation facilitates larval growth in the neotropical nymphalid butterfly Chlosyne janais. Ecological Entomology, 22(2), 133–141. https://doi.org/ https://doi.org/10.1046/j.1365-2311.1997.t01-1-00063.x

Dreyer, H., & Baumgärtner, J. (1996). Temperature influence on cohort parameters and demographic characteristics of the two cowpea coreids Clavigralla tomentosicollis and C. shadabi. Entomologia Experimentalis et Applicata, 78(2), 201–213. https://doi.org/ https://doi.org/10.1111/j.1570-7458.1996.tb00783.x

EPPO. (2004). EPPO Standards: Thaumetopoea pityocampa-PM7/37. Bulletin OEPP/EPPO Bulletin, 34, 295–298.

Fischer, E. M., & Schär, C. (2010). Consistent geographical patterns of changes in high-impact European heatwaves. Nature Geoscience, 3(6), 398–403. https://doi.org/10.1038/ngeo866

Fontaine, B., Janicot, S., & Monerie, P.-A. (2013). Recent changes in air temperature, heat waves occurrences, and atmospheric circulation in Northern Africa. Journal of Geophysical Research: Atmospheres, 118(15), 8536–8552. https://doi.org/ https://doi.org/10.1002/jgrd.50667

Forrest, J. R. K. (2016). Complex responses of insect phenology to climate change. Current Opinion in Insect Science, 17, 49–54. https://doi.org/ https://doi.org/10.1016/j.cois.2016.07.002

Gardner, J. L., Peters, A., Kearney, M. R., Joseph, L., & Heinsohn, R. (2011). Declining body size: a third universal response to warming? Trends in Ecology & Evolution, 26(6), 285–291. https://doi.org/ https://doi.org/10.1016/j.tree.2011.03.005

Georgiev, G., Rousselet, J., Laparie, M., Robinet, C., Georgieva, M., Zaemdzhikova, G., Roques, A., Bernard, A., Poitou, L., Buradino, M., Kerdelhue, C., Rossi, J. P., Matova, M., Boyadzhiev, P., & Mirchev, P. (2020). Comparative studies of egg parasitoids of the pine processionary moth (Thaumetopoea pityocampa, Den. & Schiff.) in historic and expansion areas in France and Bulgaria. Forestry: An International Journal of Forest Research, 94(2), 324–331. https://doi.org/10.1093/forestry/cpaa022

Ghosh, S. M., Testa, N. D., & Shingleton, A. W. (2013). Temperature-size rule is mediated by thermal plasticity of critical size in Drosophila melanogaster. Proceedings. Biological sciences, 280(1760), 20130174–20130174. https://doi.org/10.1098/rspb.2013.0174

Godefroid, M., Rocha, S., Santos, H., Paiva, M. R., Burban, C., Kerdelhué, C., Branco, M., Rasplus, J. Y., & Rossi, J. P. (2016). Climate constrains range expansion of an allochronic population of the pine processionary moth. Diversity and Distributions, 22(12), 1288–1300. https://doi.org/ https://doi.org/10.1111/ddi.12494

Groot, A. T., & Zizzari, Z. V. (2019). Does climate warming influence sexual chemical signaling? Animal Biology, 69(1), 83–93. https://doi.org/ https://doi.org/10.1163/15707563-20191103

GTZ, MARH, & Exaconsult Gopa. (2007). Stratégie nationale d’adaptation de l’agriculture tunisienne et des écosystèmes aux changements climatiques. Rapport d’étude dans le cadre de la coopération Tuniso-allemande publié par Deutsche Gesellschaft für Internationale.

Hickling, R., Roy, D. B., Hill, J. K., & Thomas, C. D. (2005). A northward shift of range margins in British Odonata. Global Change Biology, 11(3), 502–506. https://doi.org/ https://doi.org/10.1111/j.1365-2486.2005.00904.x

Higashi, C. H. V., Barton, B. T., & Oliver, K. M. (2020). Warmer nights offer no respite for a defensive mutualism. Journal of Animal Ecology, 89(8), 1895–1905. https://doi.org/ https://doi.org/10.1111/1365-2656.13238

Hódar, J. A., Cayuela, L., Heras, D., Pérez-Luque, A. J., & Torres-Muros, L. (2021). Expansion of elevational range in a forest pest: Can parasitoids track their hosts? Ecosphere, 12(4), e03476. https://doi.org/ https://doi.org/10.1002/ecs2.3476

Hoffmann, A. A., Anderson, A., & Hallas, R. (2002). Opposing clines for high and low temperature resistance in Drosophila melanogaster. Ecology Letters, 5(5), 614–618. https://doi.org/ https://doi.org/10.1046/j.1461-0248.2002.00367.x

Huang, Z., Ren, S., & Musa, P. D. (2008). Effects of temperature on development, survival, longevity, and fecundity of the Bemisia tabaci Gennadius (Homoptera: Aleyrodidae) predator, Axinoscymnus cardilobus (Coleoptera: Coccinellidae). Biological Control, 46(2), 209–215. https://doi.org/ https://doi.org/10.1016/j.biocontrol.2008.04.004

Huchon, H., & Demolin, G. (1970). La bioécologie de la Processionnaire du pin: dispersion potentielle, dispersion actuelle.

Imbert, C. E. (2012). Expansion d’un ravageur forestier sous l’effet du réchauffement climatique: la processionnaire du pin affecte-t-elle la biodiversité entomologique dans les zones nouvellement colonisées ? PhD dissertation, Université d’Orléans (France), pp. 198.

Jacquet, J.-S., Bosc, A., O’Grady, A. P., & Jactel, H. (2013). Pine growth response to processionary moth defoliation across a 40-year chronosequence. Forest Ecology and Management, 293, 29–38. https://doi.org/ https://doi.org/10.1016/j.foreco.2012.12.003

Jactel, H., Koricheva, J., & Castagneyrol, B. (2019). Responses of forest insect pests to climate change: not so simple. Current Opinion in Insect Science, 35, 103–108. https://doi.org/ https://doi.org/10.1016/j.cois.2019.07.010

Jeffs, C. T., & Lewis, O. T. (2013). Effects of climate warming on host–parasitoid interactions. Ecological Entomology, 38(3), 209–218. https://doi.org/ https://doi.org/10.1111/een.12026

Jones, P. D., Lister, D. H., Osborn, T. J., Harpham, C., Salmon, M., & Morice, C. P. (2012). Hemispheric and large-scale land-surface air temperature variations: An extensive revision and an update to 2010. Journal of Geophysical Research: Atmospheres, 117(D5). https://doi.org/ https://doi.org/10.1029/2011JD017139

Karban, R., & Strauss, S. Y. (2004). Physiological tolerance, climate change, and a northward range shift in the spittlebug, Philaenus spumarius. Ecological Entomology, 29(2), 251–254. https://doi.org/ https://doi.org/10.1111/j.1365-2311.2004.00576.x

Kellermann, V., Overgaard, J., Hoffmann, A. A., Fløjgaard, C., Svenning, J.-C., & Loeschcke, V. (2012). Upper thermal limits of Drosophila are linked to species distributions and strongly constrained phylogenetically. Proceedings of the National Academy of Sciences, 109(40), 16228. https://doi.org/10.1073/pnas.1207553109

Kingsolver, J. G., Diamond, S. E., & Buckley, L. B. (2013). Heat stress and the fitness consequences of climate change for terrestrial ectotherms. Functional Ecology, 27(6), 1415–1423. https://doi.org/ https://doi.org/10.1111/1365-2435.12145

Klockmann, M., Kleinschmidt, F., & Fischer, K. (2017). Carried over: Heat stress in the egg stage reduces subsequent performance in a butterfly. PloS one, 12(7), e0180968–e0180968. https://doi.org/10.1371/journal.pone.0180968

Le Lann, C., van Baaren, J., & Visser, B. (2021). Dealing with predictable and unpredictable temperatures in a climate change context: the case of parasitoids and their hosts. Journal of Experimental Biology, 224(Suppl_1). https://doi.org/10.1242/jeb.238626

Linn, C. E., Campbell, M. G., & Roelofs, W. L. (1988). Temperature modulation of behavioural thresholds controlling male moth sex pheromone response specificity. Physiological Entomology, 18(1), 59–67. https://doi.org/ https://doi.org/10.1111/j.1365-3032.1988.tb00909.x

Liu, S. S., Zhang, G. M., & Zhu, J. (1995). Influence of Temperature Variations on Rate of Development in Insects: Analysis of Case Studies from Entomological Literature. Annals of the Entomological Society of America, 88(2), 107–119. https://doi.org/10.1093/aesa/88.2.107

Martin, J. (2005). La processionnaire du pin Thaumetopoea pityocampa (Denis et Schiffermüller). Biologie et protection des forêts. Avignon: Avignon Editions, INRA, 1–62. http://www.prodinra.inra.fr/prodinra/pinra/index.xsp

Milani, N. (1990). The temperature of the egg masses of Thaumetopoea pityocampa (Den. & Schiff.) (Lepidoptera, Thaumetopoeidae). Redia, 73(1), 149–161.

Murdock, C. C., Paaijmans, K. P., Cox-Foster, D., Read, A. F., & Thomas, M. B. (2012). Rethinking vector immunology: the role of environmental temperature in shaping resistance. Nature Reviews Microbiology, 10(12), 869–876. https://doi.org/10.1038/nrmicro2900

Nangombe, S. S., Zhou, T., Zhang, W., Zou, L., & Li, D. (2019). High-Temperature Extreme Events Over Africa Under 1.5 and 2 °C of Global Warming. Journal of Geophysical Research: Atmospheres, 124(8), 4413–4428. https://doi.org/ https://doi.org/10.1029/2018JD029747

Parmesan, C., Ryrholm, N., Stefanescu, C., Hill, J. K., Thomas, C. D., Descimon, H., Huntley, B., Kaila, L., Kullberg, J., Tammaru, T., Tennent, W. J., Thomas, J. A., & Warren, M. (1999). Poleward shifts in geographical ranges of butterfly species associated with regional warming. Nature, 399(6736), 579–583. https://doi.org/ https://doi.org/10.1038/21181

Parmesan, C., & Yohe, G. (2003). A globally coherent fingerprint of climate change impacts across natural systems. Nature, 421(6918), 37–42. https://doi.org/ https://doi.org/10.1038/nature01286

Pérez-Contreras, T., & Soler, J. J. (2004). Egg parasitoids select for large clutch sizes and covering layers in pine processionary moths (Thaumetopoea pityocampa). Annales Zoologici Fennici, 41(4), 587–597. http://www.jstor.org/stable/23735942

Pérez.Contreras, T., Soler, J., & Soler, M. (2003). Why do pine processionary caterpillars Thaumetopoea pityocampa (Lepidoptera, Thaumetopoeidae) live in large groups? An experimental study. Annales Zoologici Fennici, 40, 505–515.

Pigliucci, M. (2001). Phenotypic plasticity: beyond nature and nurture. The John Hopkins University Press.

Pigliucci, M. (2005). Evolution of phenotypic plasticity: where are we going now? Trends in Ecology & Evolution, 20(9), 481–486. https://doi.org/ https://doi.org/10.1016/j.tree.2005.06.001

Pimentel, C., Ferreira, C., & Nilsson, J. A. N. Å. (2010). Latitudinal gradients and the shaping of lifehistory traits in a gregarious caterpillar. Biological Journal of the Linnean Society, 100(1), 224–236. https://doi.org/ https://doi.org/10.1111/j.1095-8312.2010.01413.x

Pincebourde, S., Dillon, M. E., & Woods, H. A. (2021). Body size determines the thermal coupling between insects and plant surfaces. Functional Ecology, 35(7), 1424–1436. https://doi.org/ https://doi.org/10.1111/1365-2435.13801

Pincebourde, S., & Woods, H. A. (2020). There is plenty of room at the bottom: microclimates drive insect vulnerability to climate change. Current Opinion in Insect Science, 41, 63–70. https://doi.org/ https://doi.org/10.1016/j.cois.2020.07.001

Poitou, L. (2021). Etude de l’impact du changement climatique sur la phénologie de la processionnaire du pin. PhD dissertation, Université d’Orléans (France), pp. 300.

Poitou, L., Laparie, M., Pincebourde, S., Rousselet, J., Suppo, C., & Robinet, C. (2022). Warming Causes Atypical Phenology in a Univoltine Moth With Differentially Sensitive Larval Stages [Original Research]. Frontiers in Ecology and Evolution, 10. https://doi.org/10.3389/fevo.2022.825875

Pureswaran, D. S., Roques, A., & Battisti, A. (2018). Forest Insects and Climate Change. Current Forestry Reports, 4(2), 35–50. https://doi.org/10.1007/s40725-018-0075-6

Quade, D. (1967). Rank Analysis of Covariance. Journal of the American Statistical Association, 62(320), 1187–1200. https://doi.org/10.2307/2283769

Raffa, K. F., Aukema, B. H., Bentz, B. J., Carroll, A. L., Hicke, J. A., Turner, M. G., & Romme, W. H. (2008). Cross-scale Drivers of Natural Disturbances Prone to Anthropogenic Amplification: The Dynamics of Bark Beetle Eruptions. BioScience, 58(6), 501–517. https://doi.org/ https://doi.org/10.1641/B580607

Ramsfield, T. D., Bentz, B. J., Faccoli, M., Jactel, H., & Brockerhoff, E. G. (2016). Forest health in a changing world: effects of globalization and climate change on forest insect and pathogen impacts. Forestry: An International Journal of Forest Research, 89(3), 245–252. https://doi.org/10.1093/forestry/cpw018

Reed, D. A., Luhring, K. A., Stafford, C. A., Hansen, A. K., Millar, J. G., Hanks, L. M., & Paine, T. D. (2007). Host defensive response against an egg parasitoid involves cellular encapsulation and melanization. Biological Control, 41(2), 214–222. https://doi.org/ https://doi.org/10.1016/j.biocontrol.2007.01.010

Reynolds, A. P., Richards, G., de la Iglesia, B., & Rayward-Smith, V. J. (2006). Clustering Rules: A Comparison of Partitioning and Hierarchical Clustering Algorithms. Journal of Mathematical Modelling and Algorithms, 5(4), 475–504. https://doi.org/10.1007/s10852-005-9022-1

Robinet, C., Baier, P., Pennerstorfer, J., Schopf, A., & Roques, A. (2007). Modelling the effects of climate change on the potential feeding activity of Thaumetopoea pityocampa (Den. & Schiff.) (Lep., Notodontidae) in France. GLOBAL ECOL BIOGEOGR, 16(4), 460–471. https://doi.org/ https://doi.org/10.1111/j.1466-8238.2006.00302.x

Robinet, C., Laparie, M., & Rousselet, J. (2015). Looking Beyond the Large Scale Effects of Global Change: Local Phenologies Can Result in Critical Heterogeneity in the Pine Processionary Moth. Frontiers in Physiology, 6(334). https://doi.org/10.3389/fphys.2015.00334

Robinet, C., & Roques, A. (2010). Direct impacts of recent climate warming on insect populations. Integrative Zoology, 5(2), 132–142. https://doi.org/ https://doi.org/10.1111/j.1749-4877.2010.00196.x

Robinet, C., Rousselet, J., Pineau, P., Miard, F., & Roques, A. (2013). Are heat waves susceptible to mitigate the expansion of a species progressing with global warming? Ecol Evol, 3(9), 2947–2957. https://doi.org/ https://doi.org/10.1002/ece3.690

Rocha, S., Kerdelhué, C., Ben Jamaa, M. L., Dhahri, S., Burban, C., & Branco, M. (2017). Effect of heat waves on embryo mortality in the pine processionary moth. Bull Entomol Res, 107(5), 583–591. https://doi.org/ https://doi.org/10.1017/S0007485317000104

Ronnås, C., Larsson, S., Pitacco, A., & Battisti, A. (2010). Effects of colony size on larval performance in a processionary moth. Ecological Entomology, 35, 436–445. https://doi.org/10.1111/j.1365-2311.2010.01199.x

Root, T. L., Price, J. T., Hall, K. R., Schneider, S. H., Rosenzweig, C., & Pounds, J. A. (2003). Fingerprints of global warming on wild animals and plants. Nature, 421(6918), 57–60. https://doi.org/10.1038/nature01333

Roques A, R. J., Avci M et al. (2015). Climate warming and past and present distribution of the processionary moths (Thaumetopoea spp.) in Europe, Asia Minor and North Africa. In R. A (Ed.), Processionary moths and climate change: an update (pp. 81–161). Springer. https://doi.org/ https://doi.org/10.1007/978-94-017-9340-7_3

Roques, L., Rossi, J.-P., Berestycki, H., Rousselet, J., Garnier, J., Roquejoffre, J.-M., Rossi, L., Soubeyrand, S., & Robinet, C. (2015). Modeling the Spatio-temporal Dynamics of the Pine Processionary Moth. In (pp. 227–263). https://doi.org/10.1007/978-94-017-9340-7_5

Rosenblatt, A. E., & Schmitz, O. J. (2016). Climate Change, Nutrition, and Bottom-Up and Top-Down Food Web Processes. Trends in Ecology & Evolution, 31(12), 965–975. https://doi.org/ https://doi.org/10.1016/j.tree.2016.09.009

Rosenzweig, C., Casassa, G., Karoly, D., Imeson, A., Liu, C., Menzel, A., Rawlins, S., Root, T., Seguin, B., & Tryjanowski, P. (2007). Assessment of observed changes and responses in natural and managed systems.. In O. F. C. M.L. Parry, J.P. Palutikof, and P.J. van der Linden (Ed.), Climate Change 2007: Impacts, Adaptation and Vulnerability. Contribution of Working Group II to the Fourth Assessment Report of the Intergovernmental Panel on Climate Change (pp. 79–131). Cambridge University Press.

Rozenberg, P., Pâques, L., Huard, F., & Roques, A. (2020). Direct and Indirect Analysis of the Elevational Shift of Larch Budmoth Outbreaks Along an Elevation Gradient [Original Research]. Frontiers in Forests and Global Change, 3(86). https://doi.org/ https://doi.org/10.3389/ffgc.2020.00086

Sales, K., Vasudeva, R., Dickinson, M. E., Godwin, J. L., Lumley, A. J., Michalczyk, Ł., Hebberecht, L., Thomas, P., Franco, A., & Gage, M. J. G. (2018). Experimental heatwaves compromise sperm function and cause transgenerational damage in a model insect. Nature Communications, 9(1), 4771. https://doi.org/10.1038/s41467-018-07273-z

Salman, M. H. R., Bonsignore, C. P., El Alaoui El Fels, A., Giomi, F., Hodar, J. A., Laparie, M., Marini, L., Merel, C., Zalucki, M. P., Zamoum, M., & Battisti, A. (2019). Winter temperature predicts prolonged diapause in pine processionary moth species across their geographic range. PeerJ, 7, e6530–e6530. https://doi.org/10.7717/peerj.6530

Santos, H., Paiva, M. R., Tavares, C., Kerdelhué, C., & Branco, M. (2011). Temperature niche shift observed in a Lepidoptera population under allochronic divergence. J. Evol. Biol, 24(9), 1897–1905. https://doi.org/ https://doi.org/10.1111/j.1420-9101.2011.02318.x

Sbay, H., & Zas, R. (2018). Geographic variation in growth, survival, and susceptibility to the processionary moth (Thaumetopoea pityocampa Dennis & Schiff.) of Pinus halepensis Mill. and P. brutia Ten.: results from common gardens in Morocco. Annals of Forest Science, 75(3), 69. https://doi.org/10.1007/s13595-018-0746-2

Schreven, S. J. J., Frago, E., Stens, A., de Jong, P. W., & van Loon, J. J. A. (2017). Contrasting effects of heat pulses on different trophic levels, an experiment with a herbivore-parasitoid model system. PloS one, 12(4), e0176704–e0176704. https://doi.org/10.1371/journal.pone.0176704

Schubert, E., & Rousseeuw, P. J. (2019). Faster k-Medoids Clustering: Improving the PAM, CLARA, and CLARANS Algorithms. In G. Amato, C. Gennaro, V. Oria, & M. Radovanović, Similarity Search and Applications Cham.

Sheridan, J. A., & Bickford, D. (2011). Shrinking body size as an ecological response to climate change. Nature Climate Change, 1(8), 401–406. https://doi.org/10.1038/nclimate1259

Sinclair, B. J., Williams, C. M., & Terblanche, J. S. (2012). Variation in Thermal Performance among Insect Populations. Physiological and Biochemical Zoology: Ecological and Evolutionary Approaches, 85(6), 594–606. https://doi.org/10.1086/665388

Stireman, J. O., Dyer, L. A., Janzen, D. H., Singer, M. S., Lill, J. T., Marquis, R. J., Ricklefs, R. E., Gentry, G. L., Hallwachs, W., Coley, P. D., Barone, J. A., Greeney, H. F., Connahs, H., Barbosa, P., Morais, H. C., & Diniz, I. R. (2005). Climatic unpredictability and parasitism of caterpillars: Implications of global warming. Proceedings of the National Academy of Sciences of the United States of America, 102(48), 17384. https://doi.org/10.1073/pnas.0508839102

Thompson, R. M., Beardall, J., Beringer, J., Grace, M., & Sardina, P. (2013). Means and extremes: building variability into community-level climate change experiments. Ecology Letters, 16(6), 799–806. https://doi.org/ https://doi.org/10.1111/ele.12095

Vasseur, D. A., DeLong, J. P., Gilbert, B., Greig, H. S., Harley, C. D. G., McCann, K. S., Savage, V., Tunney, T. D., & O’Connor, M. I. (2014). Increased temperature variation poses a greater risk to species than climate warming. Proceedings of the Royal Society B: Biological Sciences, 281(1779), 20132612. https://doi.org/ doi:10.1098/rspb.2013.2612

Verner, D., Wilby, R., Breisinger, C., Al-Riffai, P., Robertson, R., Wiebelt, M., Kronik, J., Clement, V., Levine, T., Esen, F., & Roos, P. (2013). Tunisia in a changing climate: assessment and actions for increased resilience and development. World Bank Publications. https://doi.org/ https://doi.org/10.1596/978-0-8213-9857-9

Visser, M. E., & Holleman, L. J. M. (2001). Warmer springs disrupt the synchrony of oak and winter moth phenology. Proceedings of the Royal Society of London. Series B: Biological Sciences, 268(1464), 289–294. https://doi.org/ doi:10.1098/rspb.2000.1363

Walther, G.-R., Post, E., Convey, P., Menzel, A., Parmesan, C., Beebee, T. J. C., Fromentin, J.-M., Hoegh-Guldberg, O., & Bairlein, F. (2002). Ecological responses to recent climate change. Nature, 416(6879), 389–395. https://doi.org/ https://doi.org/10.1038/416389a

Walther, G. R. (2010). Community and ecosystem responses to recent climate change. Philosophical Transactions of the Royal Society B: Biological Sciences, 365(1549), 2019–2024. https://doi.org/ doi:10.1098/rstb.2010.0021

Wertheim, B., Kraaijeveld, A. R., Schuster, E., Blanc, E., Hopkins, M., Pletcher, S. D., Strand, M. R., Partridge, L., & Godfray, H. C. J. (2005). Genome-wide gene expression in response to parasitoid attack in Drosophila. Genome Biology, 6(11), R94. https://doi.org/10.1186/gb-2005-6-11-r94

Woods, H. A., Dillon, M. E., & Pincebourde, S. (2015). The roles of microclimatic diversity and of behavior in mediating the responses of ectotherms to climate change. Journal of Thermal Biology, 54, 86–97. https://doi.org/ https://doi.org/10.1016/j.jtherbio.2014.10.002

Wu, C.-H., Holloway, J. D., Hill, J. K., Thomas, C. D., Chen, I. C., & Ho, C.-K. (2019). Reduced body sizes in climate-impacted Borneo moth assemblages are primarily explained by range shifts. Nature Communications, 10(1), 4612. https://doi.org/10.1038/s41467-019-12655-y

Zhao, F., Zhang, W., Hoffmann, A. A., & Ma, C. S. (2014). Night warming on hot days produces novel impacts on development, survival and reproduction in a small arthropod. Journal of Animal Ecology, 83(4), 769–778. https://doi.org/ https://doi.org/10.1111/1365-2656.12196

Zittis, G., Hadjinicolaou, P., Almazroui, M., Bucchignani, E., Driouech, F., El Rhaz, K., Kurnaz, L., Nikulin, G., Ntoumos, A., Ozturk, T., Proestos, Y., Stenchikov, G., Zaaboul, R., & Lelieveld, J. (2021). Business-as-usual will lead to super and ultra-extreme heatwaves in the Middle East and North Africa. npj Climate and Atmospheric Science, 4(1), 20. https://doi.org/10.1038/s41612-021-00178-7

