## Supplementary material (11 items) for "Effects of climate warming on the pine processionary moth at the southern edge of its range: a retrospective analysis on egg survival in Tunisia"

### Supplementary information

**Table SM1.** Coordinates of sampling sites.

| Field egg masses sampling sites | Latitude (°N) | Longitude (°E) |
|---|---|---|
| Chaambi | 35.200000 | 8.700000 |
| El Ayoun | 35.557653 | 8.879097 |
| Mghila | 35.333333 | 9.200000 |
| Thélepte | 34.960611 | 8.583472 |
| Bir Lahfay | 34.944697 | 9.156361 |
| El Fej | 34.701582 | 9.043508 |
| Jebel Motlag | 34.954417 | 9.707389 |
| Jebel Rihane | 34.823708 | 9.636170 |
| cité Ettahrir | 36.821750 | 10.135778 |
| Dar Chichou | 36.965594 | 10.964017 |
| Korbous | 36.833333 | 10.583333 |
| Sejnane | 37.183333 | 9.183333 |
| Oued Laabid | 36.816542 | 10.711303 |
| El Menzeh | 36.837850 | 10.184692 |
| Manouba | 36.814722 | 10.108361 |
| Ain Jamala | 36.503839 | 9.301167 |
| Testour | 36.544944 | 9.377444 |
| Henchir Naam | 36.216667 | 9.166667 |
| Jebel Kbouch (Kef) | 36.210000 | 8.930000 |
| El Krib (Siliana) | 36.332544 | 9.063128 |
| Kesra (Siliana) | 35.871833 | 9.366583 |
| Jebel Sidi Aich (Gafsa) | 34.783292 | 8.865861 |

**Table SM2.** Temperature datasets (combination of data from the Institut National de Météorologie, INM, and the NASA data in corresponding grid cells of 0.5 degree × 0.625 degree (roughly 50 × 60 km)) and coordinates of meteorological stations.

| INM meteorological station | Latitude (°N) | Longitude (°E) | Available data (INM) | Data from NASA used to complete the INM datasets |
|---|---|---|---|---|
| Kélibia | 36.844855 | 11.082701 | 2001-2011 | 1990-2000<br>2012-2019 |
| Carthage | 36.846081 | 10.219053 | 1990-2014 | 2015-2019 |
| Béja | 36.723338 | 9.184013 | 1990-1997<br>2001-2011 | 1998-2000<br>2012-2019 |
| Siliana | 35.851853 | 9.595147 | 1990-1997 | 1998-2019 |
| Kef | 36.120862 | 8.720267 | 1990-1997<br>2001-2011 | 1998-2000<br>2012-2019 |
| Kasserine | 34.948369 | 8.569550 | 2001-2011 | 1990-2000<br>2012-1019 |
| Sidi Bouzid | 35.025685 | 9.498840 | 1990-2014 | 2015-2019 |
| Gafsa | 34.427352 | 8.820959 | 1990-2014 | 2015-2019 |

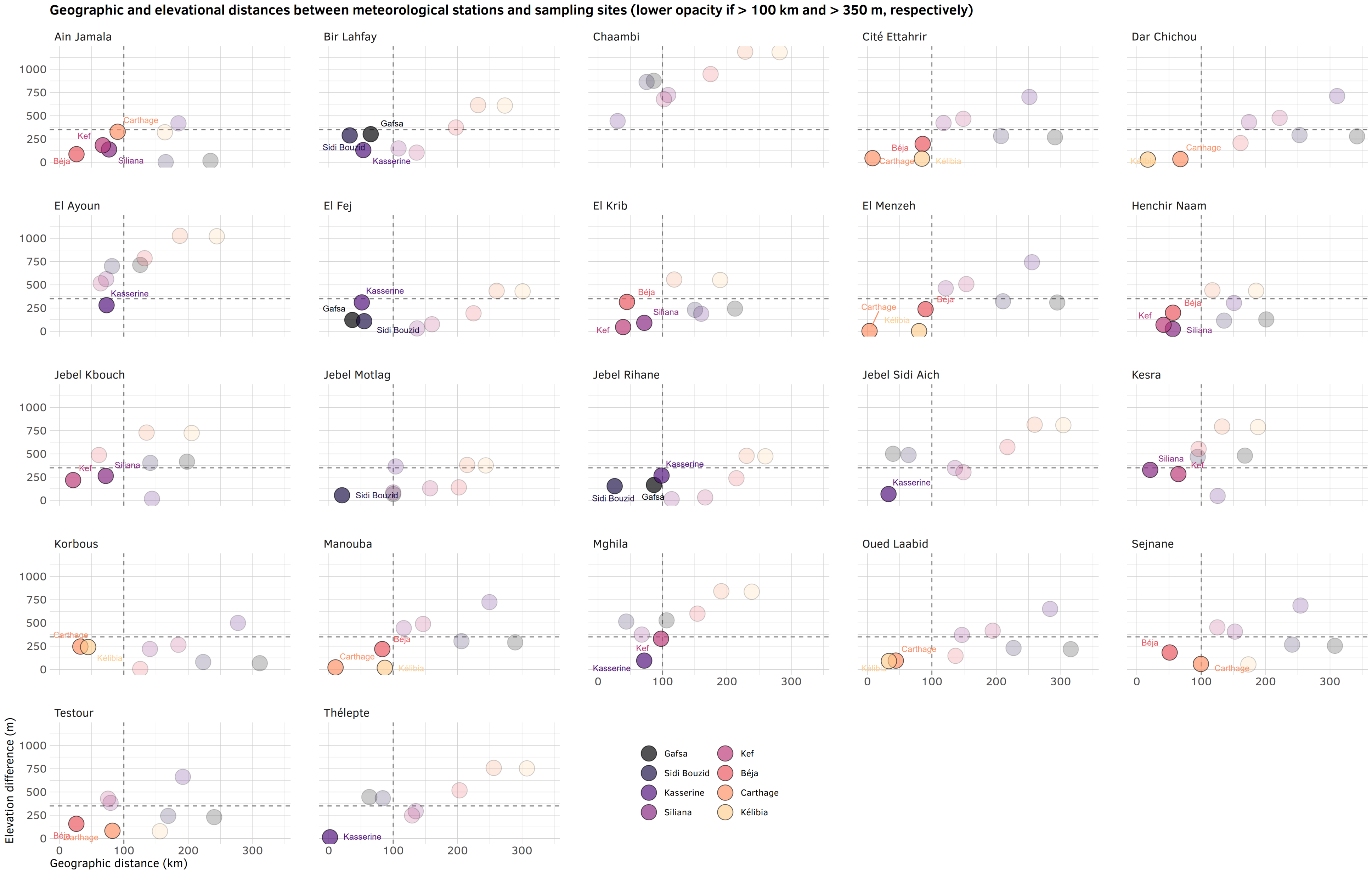


**Figure SM3.** Geographic and elevational distances between meteorological stations and egg sampling sites (lower opacity if >100 km and >350 m, respectively).

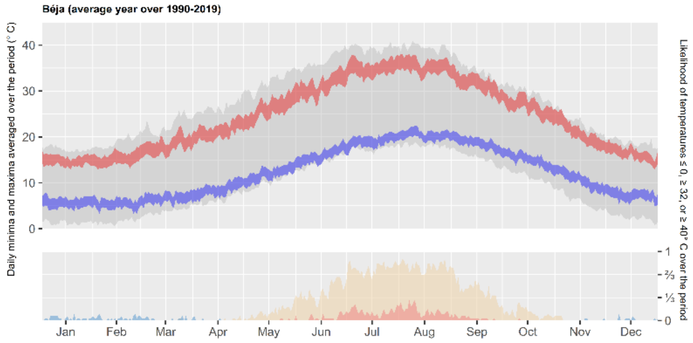
Béja (average year over 1990-2019)
Daily minima and maxima averaged over the period (° C)
Likelihood of temperatures ≤ 0, ≥ 32, or ≥ 40° C over the period
Jan Feb Mar Apr May Jun Jul Aug Sep Oct Nov Dec

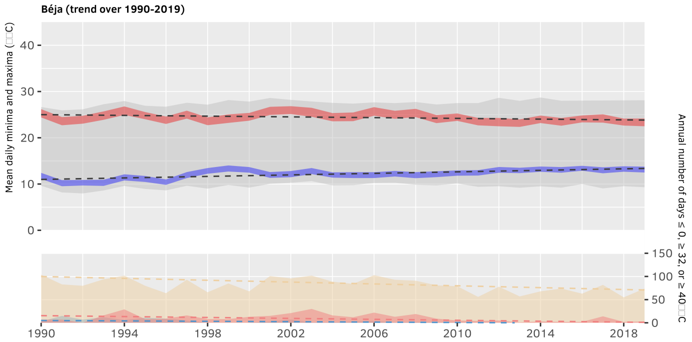
Béja (trend over 1990-2019)
Mean daily minima and maxima
Annual number of days ≤ 0, ≥ 32, or ≥ 40
1990 1994 1998 2002 2006 2010 2014 2018

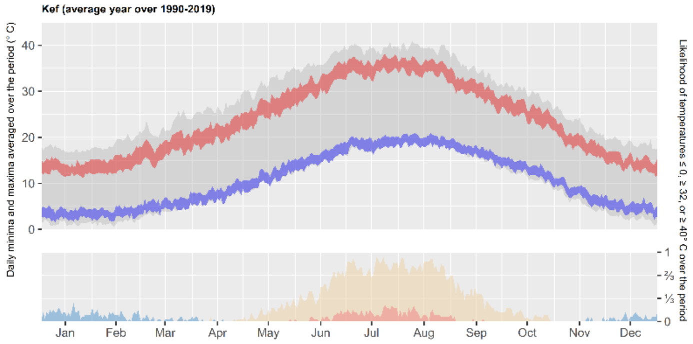
Kef (average year over 1990-2019)
Daily minima and maxima averaged over the period (° C)
Likelihood of temperatures ≤ 0, ≥ 32, or ≥ 40° C over the period
Jan Feb Mar Apr May Jun Jul Aug Sep Oct Nov Dec

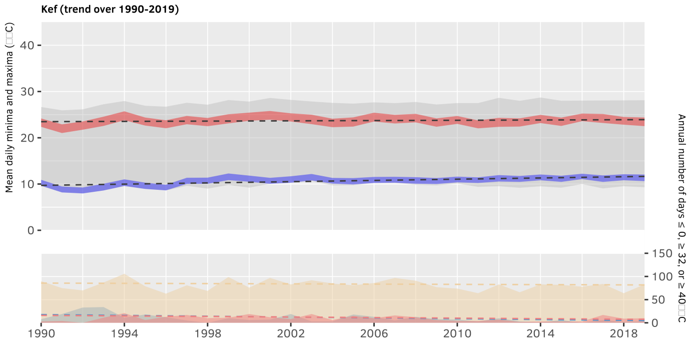
Kef (trend over 1990-2019)
Mean daily minima and maxima
Annual number of days ≤ 0, ≥ 32, or ≥ 40
1990 1994 1998 2002 2006 2010 2014 2018

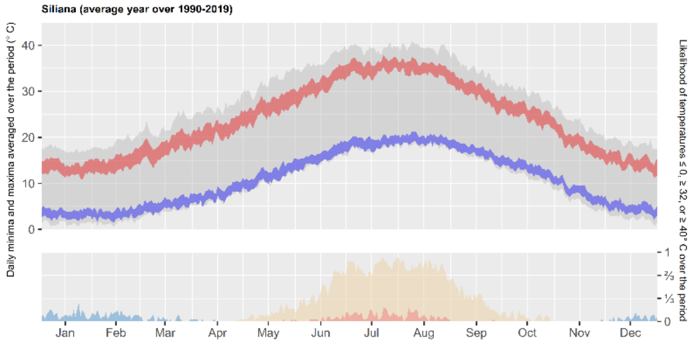
Siliana (average year over 1990-2019)
Daily minima and maxima averaged over the period (° C)
Likelihood of temperatures ≤ 0, ≥ 32, or ≥ 40° C over the period
Jan Feb Mar Apr May Jun Jul Aug Sep Oct Nov Dec

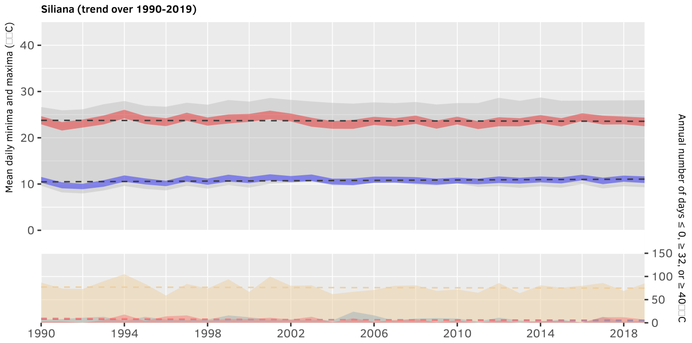
Siliana (trend over 1990-2019)
Mean daily minima and maxima
Annual number of days ≤ 0, ≥ 32, or ≥ 40
1990 1994 1998 2002 2006 2010 2014 2018

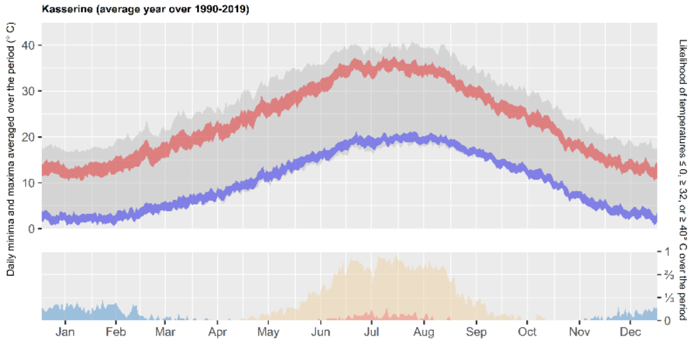
Kasserine (average year over 1990-2019)
Daily minima and maxima averaged over the period (° C)
Likelihood of temperatures ≤ 0, ≥ 32, or ≥ 40° C over the period
Jan Feb Mar Apr May Jun Jul Aug Sep Oct Nov Dec

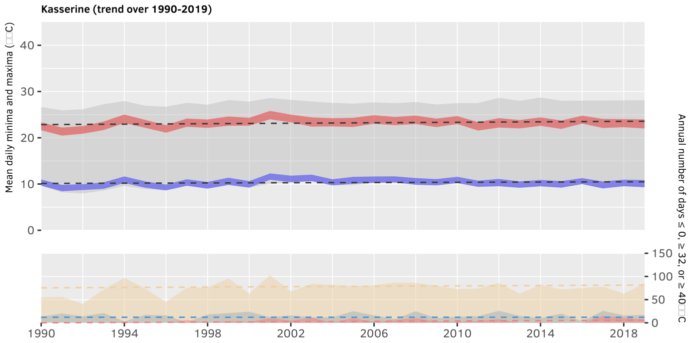
Kasserine (trend over 1990-2019)
Mean daily minima and maxima
Annual number of days ≤ 0, ≥ 32, or ≥ 40
1990 1994 1998 2002 2006 2010 2014 2018

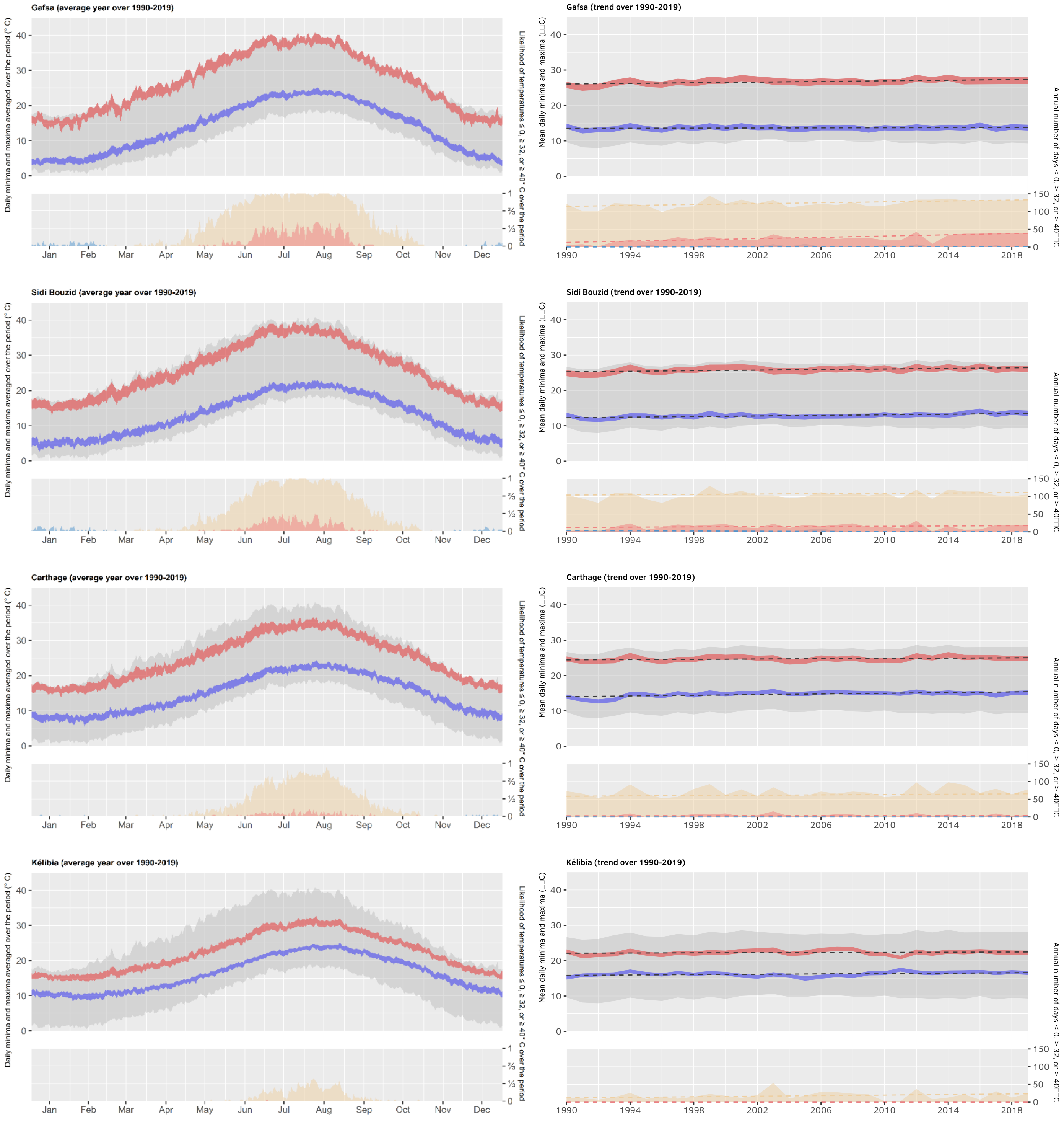


**Figure SM4.** Climate data in eight Tunisian regions between 1990 and 2019 (data source: INM and NASA, see Table SM2). Left charts show the mean year in each region by averaging daily maxima (red) and minima (blue) by day of the year over the period, represented as 95% CI ribbons. Right charts show the yearly average of daily maxima (red) and minima (blue), represented as 95% CI ribbons, and corresponding Theil-Sen estimators. Thick grey ribbons in the background show the maximal thermal range across all nine regions depending on day of the year (left) or year (right). The bottom part of left charts shows the likelihood of temperatures ≤ 0 (blue), ≥ 32 (beige) or ≥ 40 °C (red), while the bottom part of right charts shows the annual number of days below or above those thresholds. The 366$^{th}$ day during leap years was discarded due to its lower sample size.

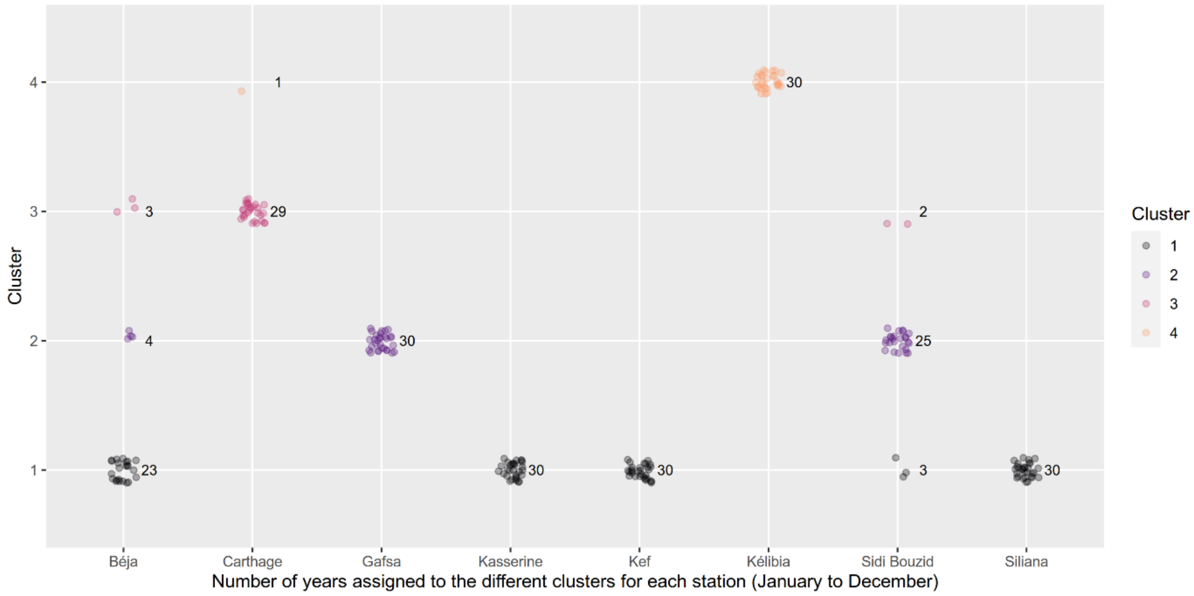


**Figure SM5.** Number of years assigned to the different clusters for each meteorological series (PAM clustering on data from January to December).

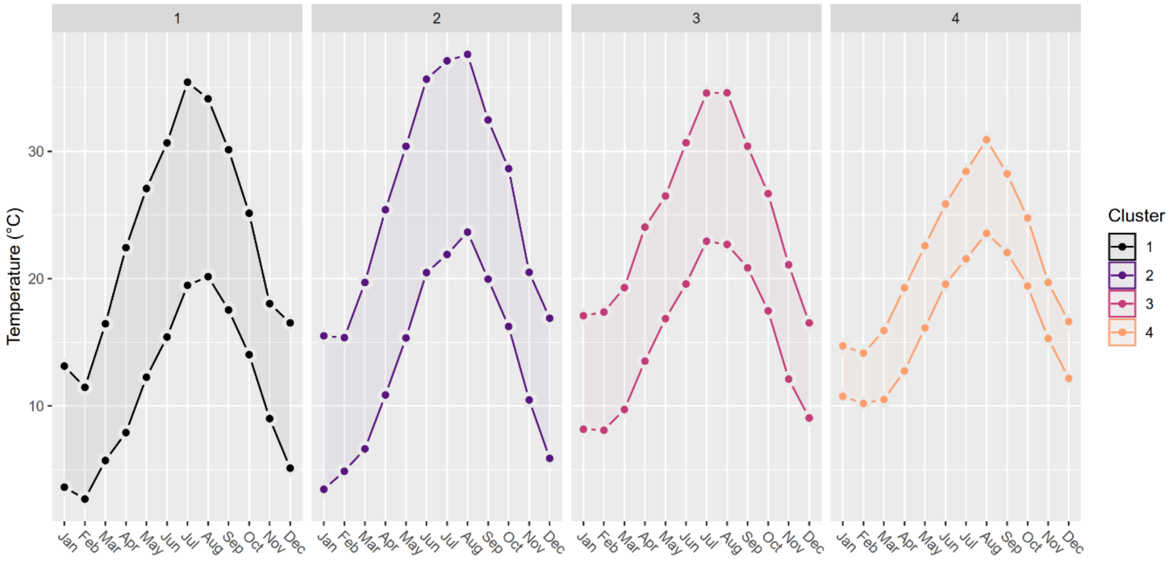


**Figure SM6.** TX and TN from January to December in each cluster medoid (PAM clustering on data from January to December).

**SM7.** Synchronic comparison of egg phenotypes among all four clusters.

**A.** Analysis with all data including cluster 1 in 2017.

A Quade's RANCOVA (see details on this method in subsection 3 of Materials and Methods) was used to compare egg phenotypes among all four clusters synchronically, *i.e.*, only within the period(s) clusters have in common (no interaction term). It was followed by a Tukey post hoc procedure to investigate differences when a significant effect was found.

***Clutch size.*** The analysis revealed no differences among clusters 1, 3 and 4 in the 1990s (Quade's RANCOVA, $F_{dt} = 1.58_2$, p = 0.208), whereas it confirmed significant differences among clusters 1, 2 and 3 in the 2010s ($F_{df} = 4.40_2$, p = 0.013). Specifically, fecundity was significantly higher in cluster 1 than cluster 3 (Tukey post hoc, p = 0.009).

***Hatching rate and egg mortality factors.*** The results showed synchronic differences among clusters in the rates of hatched, sterile, aborted and parasitized eggs (see the following two tables for details).

Results of Tukey's post hoc tests in the 1990s.

| Variable | Clusters | Mean difference (I-J) | Standard error | *p* value |
|---|---|---|---|---|
| Hatching rate | 1 vs 3 | 43.5* | 11.6 | 0.001 |
| | 1 vs 4 | -43.9* | 15 | 0.01 |
| | 3 vs 4 | -87.4* | 16.2 | <0.001 |
| Sterility rate | 1 vs 3 | 9.8 | 11.8 | 0.68 |
| | 1 vs 4 | 32.1 | 15.3 | 0.09 |
| | 3 vs 4 | 22.3 | 16.5 | 0.37 |
| Abortion rate | 1 vs 3 | -42.8* | 11.8 | 0.001 |
| | 1 vs 4 | -13.9 | 15.3 | 0.64 |
| | 3 vs 4 | 29 | 16.5 | 0.19 |
| Parasitism rate | 1 vs 3 | -29.8* | 11.4 | 0.03 |
| | 1 vs 4 | 72* | 14.8 | <0.001 |
| | 3 vs 4 | 101.4* | 16 | <0.001 |

*. The mean difference is significant at the .05 level.

Results of Tukey's post hoc tests in the 2010s.

| Variable | Clusters | Mean difference (I-J) | Standard error | *p* value |
|---|---|---|---|---|

| | | | | |
|---|---|---|---|---|
| Hatching rate | 1 vs 2 | -18.7 | 13.1 | 0.33 |
| | 1 vs 3 | -46.7* | 16.8 | 0.02 |
| | 2 vs 3 | -28 | 18.3 | 0.28 |
| Sterility rate | 1 vs 2 | 16.5 | 13.2 | 0.42 |
| | 1 vs 3 | 59.2* | 16.9 | <0.001 |
| | 2 vs 3 | 42.6 | 18.4 | 0.054 |
| Abortion rate | 1 vs 2 | -34.3* | 13.2 | 0.03 |
| | 1 vs 3 | -35.2 | 17 | 0.10 |
| | 2 vs 3 | -0.9 | 18.5 | 0.99 |
| Parasitism rate | 1 vs 2 | -21.3 | 13.1 | 0.24 |
| | 1 vs 3 | -48.1* | 16.9 | 0.01 |
| | 2 vs 3 | -26.8 | 18.4 | 0.31 |

*. The mean difference is significant at the .05 level.

**B.** Analysis without data from cluster 1 in 2017.

A Quade's RANCOVA (see details on this method in subsection 3 of Materials and Methods) was used to compare egg phenotypes among all four clusters synchronically, *i.e.*, only within the period(s) clusters have in common (no interaction term), excluding data from cluster 1 in 2017. It was followed by a Tukey post hoc procedure to investigate differences when a significant effect was found.

The analysis revealed patterns different from those found when considering data from cluster 1 in 2017, since no difference among clusters appeared in the 2010s in any of the variables (Quade's RANCOVA, hatching rate: $F_{df} = 0.9_2$, p = 0.412; sterility rate: $F_{df} = 2.8_2$, p = 0.063; abortion rate: $F_{df} = 0.01_2$, p = 0.993; parasitism rate: $F_{df} = 1.5_2$, p = 0.218).

**Table SM8.** Descriptive statistics with all data including cluster 1 in 2017: observed mean (M), Quade's adjusted mean (Madj) and associated standard error (SE) for the different response variables.

| Variable | Period | Cluster | M (SE) | Madj |
|---|---|---|---|---|
| Fecundity | 1990s | 1 | 156.6 (3) | -0.2(6.9) |
| | | 2 | NA | NA |
| | | 3 | 159.7 (4.3) | 10 (10.2) |
| | | 4 | 145.4(6.3) | -20 (14.5) |
| | 2010s | 1 | 153 (3) | 10.7 (7.8) |
| | | 2 | 142.1 (4.5) | -0.3 (10.8) |
| | | 3 | 134.5 (5.6) | -39.7 (14.7) |
| | | 4 | NA | NA |
| Hatching rate | 1990s | 1 | 73.5 (1.7) | 6.9 (7.1) |
| | | 2 | NA | NA |
| | | 3 | 63.3 (2.5) | -36.6 (8.7) |
| | | 4 | 79.3 (3.1) | 50.8 (14.1) |
| | 2010s | 1 | 46.8 (2.5) | -12.4 (8) |
| | | 2 | 48.9 (3.4) | 6.2 (10.6) |
| | | 3 | 63.2 (3.6) | 34.2 (12.6) |
| | | 4 | NA | NA |
| Sterility rate | 1990s | 1 | 6.8 (0.6) | 7.8 (7.3) |
| | | 2 | NA | NA |
| | | 3 | 6.6 (0.7) | -2 (9.3) |
| | | 4 | 5.8 (0.7) | -24.3 (12.8) |
| | 2010s | 1 | 27.1 (2.5) | 13.6 (8.4) |
| | | 2 | 17.3 (2.6) | -2.9 (9.9) |
| | | 3 | 6.6 (1.7) | -45.5 (11.7) |
| | | 4 | NA | NA |
| Abortion rate | 1990s | 1 | 2.4 (0.2) | -15.3 (6.6) |
| | | 2 | NA | NA |
| | | 3 | 9.3 (1.9) | 27.5 (9.9) |
| | | 4 | 8.1 (2.8) | -1.4 (15.4) |
| | 2010s | 1 | 17.2 (1.8) | -15.4 (8) |
| | | 2 | 24.4 (2.9) | 18.9 (10.7) |
| | | 3 | 17.4 (2.9) | 19.8 (12.8) |
| | | 4 | NA | NA |
| Parasitism rate | 1990s | 1 | 17.2 (1.3) | 1.4 (6.8) |
| | | 2 | NA | NA |
| | | 3 | 20.8 (1.7) | 31.2 (9.4) |
| | | 4 | 6.8 (1.5) | -70.1 (13) |
| | 2010s | 1 | 8.9 (0.7) | -13.4 (8.3) |
| | | 2 | 9.4 (1.1) | 7.9 (9.8) |
| | | 3 | 12.8 (1.8) | 34.7 (13.3) |
| | | 4 | NA | NA |

**Table SM9.** Descriptive statistics without data from cluster 1 in 2017: observed mean (M), Quade's adjusted mean (Madj) and associated standard error (SE) for the different response variables.

| Variable | Period | Cluster | M (SE) | Madj (SE) |
|---|---|---|---|---|
| Fecundity | 1990s | 1 | 156.6 (3) | -0.2(6.9) |
| | | 2 | NA | NA |
| | | 3 | 159.7 (4.3) | 10 (10.2) |
| | | 4 | 145.4 (6.3) | -20 (14.5) |
| | 2010s | 1 | 155.9 (3.3) | 13.6 (7.7) |
| | | 2 | 142.1 (4.5) | -2.5 (9.6) |
| | | 3 | 134.5 (5.6) | -36.6 (13.2) |
| | | 4 | NA | NA |
| Hatching rate | 1990s | 1 | 73.5 (1.7) | 6.9 (7.1) |
| | | 2 | NA | NA |
| | | 3 | 63.3 (2.5) | -36.6 (8.7) |
| | | 4 | 79.3 (3.1) | 50.8 (14.1) |
| | 2010s | 1 | 57.6 (2.5) | 1.3 (7.8) |
| | | 2 | 48.9 (3.4) | -8.6 (9.9) |
| | | 3 | 63.2 (3.6) | 13.2 (12.2) |
| | | 4 | NA | NA |
| Sterility rate | 1990s | 1 | 6.8 (0.6) | 7.8 (7.3) |
| | | 2 | NA | NA |
| | | 3 | 6.6 (0.7) | -2 (9.3) |
| | | 4 | 5.8 (0.7) | -24.3 (12.8) |
| | 2010s | 1 | 10.2 (1.1) | -2.5 (8) |
| | | 2 | 17.3 (2.6) | 15.4 (9.7) |
| | | 3 | 6.6 (1.7) | -23.1 (11.7) |
| | | 4 | NA | NA |
| Abortion rate | 1990s | 1 | 2.4 (0.2) | -15.3 (6.6) |
| | | 2 | NA | NA |
| | | 3 | 9.3 (1.9) | 27.5 (9.9) |
| | | 4 | 8.1 (2.8) | -1.4 (15.4) |
| | 2010s | 1 | 21.2 (2.1) | -0.6 (7.8) |
| | | 2 | 24.4 (2.9) | 0.8 (10.3) |
| | | 3 | 17.4 (2.9) | 0.5 (12.3) |
| | | 4 | NA | NA |
| Parasitism rate | 1990s | 1 | 17.2 (1.3) | 1.4 (6.8) |
| | | 2 | NA | NA |
| | | 3 | 20.8 (1.7) | 31.2 (9.4) |
| | | 4 | 6.8 (1.5) | -70.1 (13) |
| | 2010s | 1 | 11 (0.8) | 3 (8.1) |
| | | 2 | 9.4 (1.1) | -12.3 (9.3) |
| | | 3 | 12.8 (1.8) | 15.4 (12.9) |
| | | 4 | NA | NA |

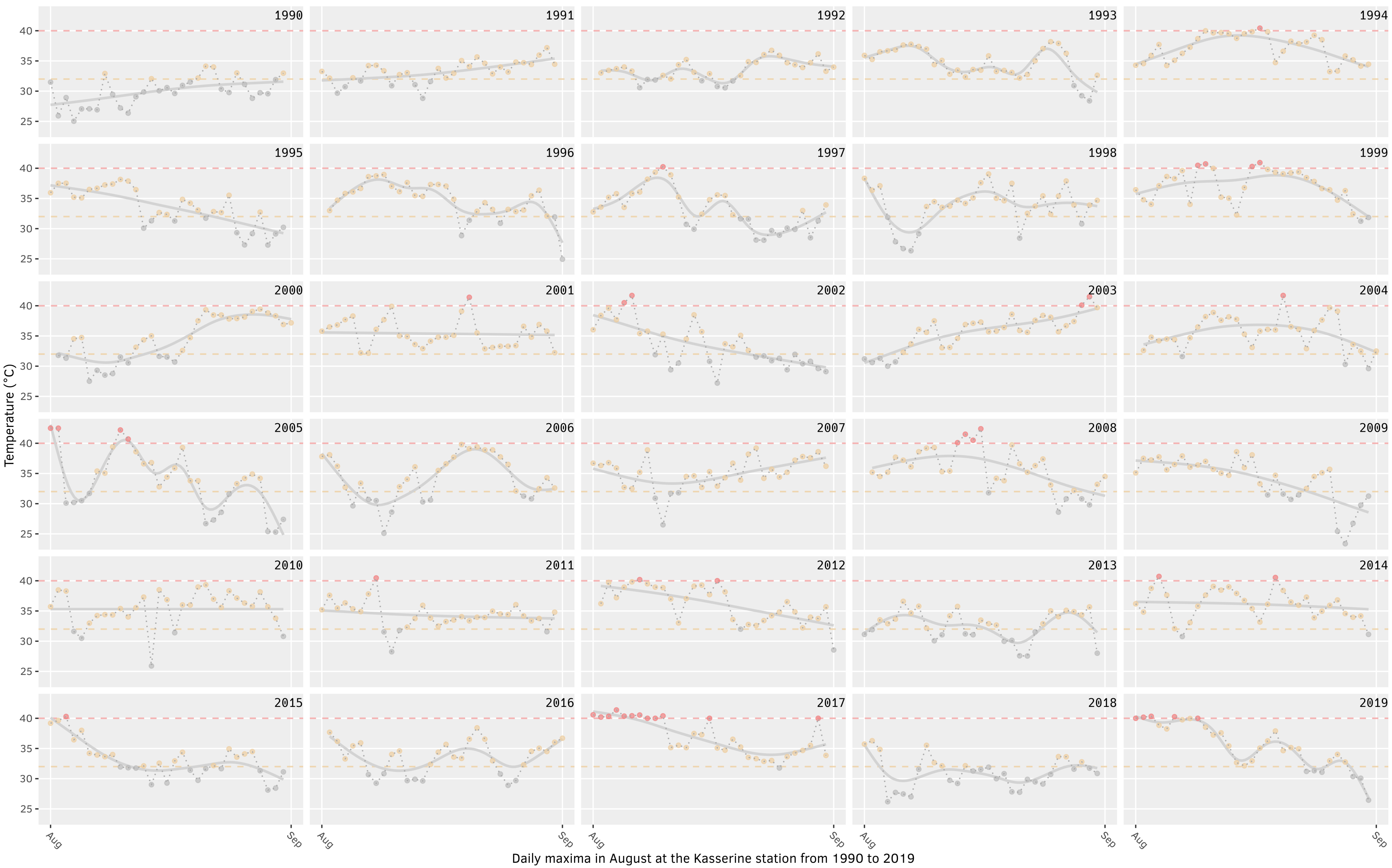


**Figure SM10.** August daily maximal temperature recorded in 1990-2019 at the Kasserine station, near the sampling site of Thélepte (cluster 1). Yellow points correspond to daily maxima ≥ 32 °C, red points correspond to daily maxima ≥ 40 °C. Smooth lines are fitted with the "gam" (generalized additive model) method.

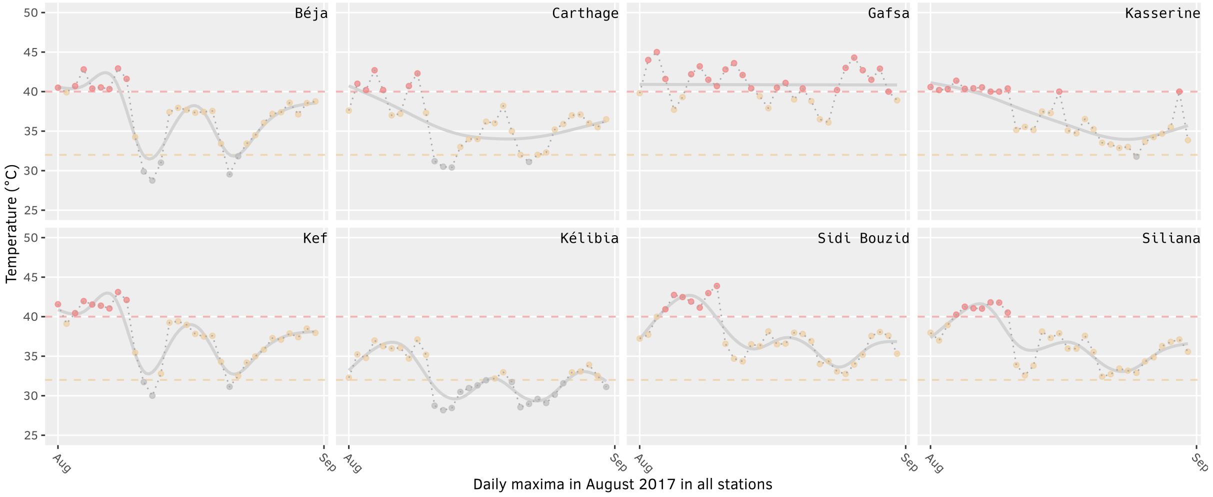


**Figure SM11.** Daily maximal temperature recorded in August 2017 in all stations. Béja, Kasserine, Kef and Siliana belong to Cluster 1; Gafsa and Sidi Bouzid belong to Cluster 2; Carthage belongs to Cluster 3; Kélibia belongs to Cluster 4. Yellow points correspond to daily maxima ≥ 32 °C, red points correspond to daily maxima ≥ 40 °C. Smooth lines are fitted with the “gam” (generalized additive model) method.
